# Testing the inferred transcription rates of a dynamic, gene network model in absolute units

**DOI:** 10.1101/2021.03.18.436071

**Authors:** Uriel Urquiza-García, Andrew J. Millar

## Abstract

The circadian clock coordinates plant physiology and development. Mathematical clock models have provided a rigorous framework to understand how the observed rhythms emerge from disparate, molecular processes. However, models of the plant clock have largely been built and tested against RNA timeseries data in arbitrary, relative units. This limits model transferability, refinement from biochemical data and applications in synthetic biology. Here, we incorporate absolute mass units into a detailed, gene circuit model of the clock in *Arabidopsis thaliana*. We re-interpret the established P2011 model, highlighting a transcriptional activator that overlaps the function of REVEILLE 8/LHY-CCA1-LIKE 5, and refactor dynamic equations for the Evening Complex. The U2020 model incorporates the repressive regulation of *PRR* genes, a key feature of the most detailed clock model F2014, without greatly increasing model complexity. We tested the experimental error distributions of qRT-PCR data calibrated for units of RNA transcripts/cell and of circadian period estimates, in order to link the models to data more appropriately. U2019 and U2020 models were constrained using these data types, recreating previously-described circadian behaviours with RNA metabolic processes in absolute units. To test their inferred rates, we estimated a distribution of observed, transcriptome-wide transcription rates (Plant Empirical Transcription Rates, PETR) in units of transcripts/cell/hour. The PETR distribution and the equivalent degradation rates indicated that the models’ predicted rates are biologically plausible, with individual exceptions. In addition to updated, explanatory models of the plant clock, this validation process represents an advance in biochemical realism for models of plant gene regulation.

## Introduction

The circadian clock in plants temporally coordinates an extensive repertoire of developmental and physiological processes. These include seedling establishment, photosynthesis, cell division, and flowering time, amongst others. Its physiological relevance, architecture and response to environmental signals have been extensively reviewed (Millar 2016; Nohales and Kay 2016; Sanchez *et al*. 2020). The fact that mutations of the clock genes have such pleiotropic effects can be explained at the molecular level, by observations that more than 30% of the *Arabidopsis thaliana* transcriptome exhibits circadian rhythmicity (Covington *et al*. 2008; Edwards *et al*. 2006; Harmer *et al*. 2000). The seminal timeseries studies have been repeated in other species, demonstrating how the Arabidopsis results translate into crops (Edwards *et al*. 2018; Li *et al*. 2019; Matsuzaki *et al*. 2015; L. M. Müller *et al*. 2020; N. A. Müller *et al*. 2016). Studies of seasonal timing (phenology) in crops are also uncovering the effects of clock genes that are homologous to those first identified in Arabidopsis, and the impact of Genotype x Environment interactions (Bendix *et al*. 2015; Millar 2016). Understanding the clock gene circuit and its outputs to physiology and phenology therefore holds promise to guide further breeding and/or engineering of crop varieties (Preuss *et al*. 2012 p. 32) and the clock regulation of RNA levels is of particular interest.

*Arabidopsis thaliana* remains a critical, model organism for prototyping the theoretical and experimental tools that are subsequently combined with the methods from crop science. Systems biology approaches have long been applied to understand the complex mechanism of the Arabidopsis clock gene circuit, as noted in the reviews cited above, and introduced in detail below. However, even this paradigmatic system still falls short of the resources that are required to understand (explain and predict, quantitatively and mechanistically) how genome sequence variation at single base-pair resolution affects whole-organism traits (Millar 2016). This also hampers the selection of the genome sequence that is needed for some desired physiology (Marshall-Colon *et al*. 2017) whereas this possibility is already exemplified in other systems, such as the engineering of transcription factor binding sites in *E. coli* (Barnes *et al*. 2019). Aspects of the plant clock mechanism based on RNA metabolism have been more tractable than protein regulation to date, whether in RNA processing (James *et al*. 2012) or as here in transcriptional regulation (Kang *et al*. 2018; Lin *et al*. 2020; Xu *et al*. 2020). Even at this level, current models of plant gene regulatory networks represent their RNA components with arbitrary mass units, and hence their transcription and translation rates likewise. This practice is inherited from the experimental methodology, where RNA abundance is typically normalised to an internal standard for quantification (Czechowski *et al*. 2005), yielding data in arbitrary, relative units.

Refactoring the models to use absolute mass units would not only be a step towards engineering but should also allow more discriminating tests of model validity. First, any estimated parameter value should fall within a realistic distribution for the cognate biochemical process. Such a distribution is hard to generalise in arbitrary units. This test can already discriminate between models and reveal the system’s evolutionary constraints, as shown for enzymatic reactions (Bar-Even *et al*. 2011). Collating biochemical parameter measurements to form an expected distribution is therefore beneficial (Jolley *et al*. 2014), similarly to the collation of trait data for physiology (Poorter *et al*. 2010), but has been rare for plant gene regulation (Millar *et al*. 2019). Here, we aim to contribute at this level. Second and more obviously, biochemical measurements should directly test the particular parameter values in the model, improving the model’s realism over time.

This approach is most relevant to models that focus on biochemical fidelity (Foo *et al*. 2016; J. C. W. Locke *et al*. 2005; James C. W. Locke *et al*. 2005; Pokhilko *et al*. 2010, 2012, 2013; Zeilinger *et al*. 2006). Other clock models have used greater levels of abstraction to emphasise operating principles (De Caluwé *et al*. 2016, 2017; Foo *et al*. 2016, 2020) or to gain computational tractability when studying large ensembles (Greenwood *et al*. 2020). Our interest in linking genome sequence mechanistically to complex plant phenotypes requires significant biochemical and biophysical detail, so we consider two of the most detailed models available, referred to as P2011 (Pokhilko *et al*. 2012) and F2014 (Fogelmark and Troein 2014).

### The clock mechanism represented in P2011 and F2014

Both models represent a set of interlocking, transcriptional, negative feedback loops, in which the clock genes control each other’s expression at particular times of day, through their short-lived proteins (Figure 1a; Supplementary Table 1). 24-hour rhythms of gene activity with characteristic waveforms emerge from the gene network models’ dynamics, without a single ‘rhythm generator’. Clock gene expression is affected by several light input pathways, so the biological rhythms are entrained to the day/night cycle. Both models reflect the rhythmic expression profiles of clock components observed in several light conditions and mutant backgrounds with good fidelity, and in real time units, though they differ in several, relevant details (please see Supplementary Information).

**Figure 1.**
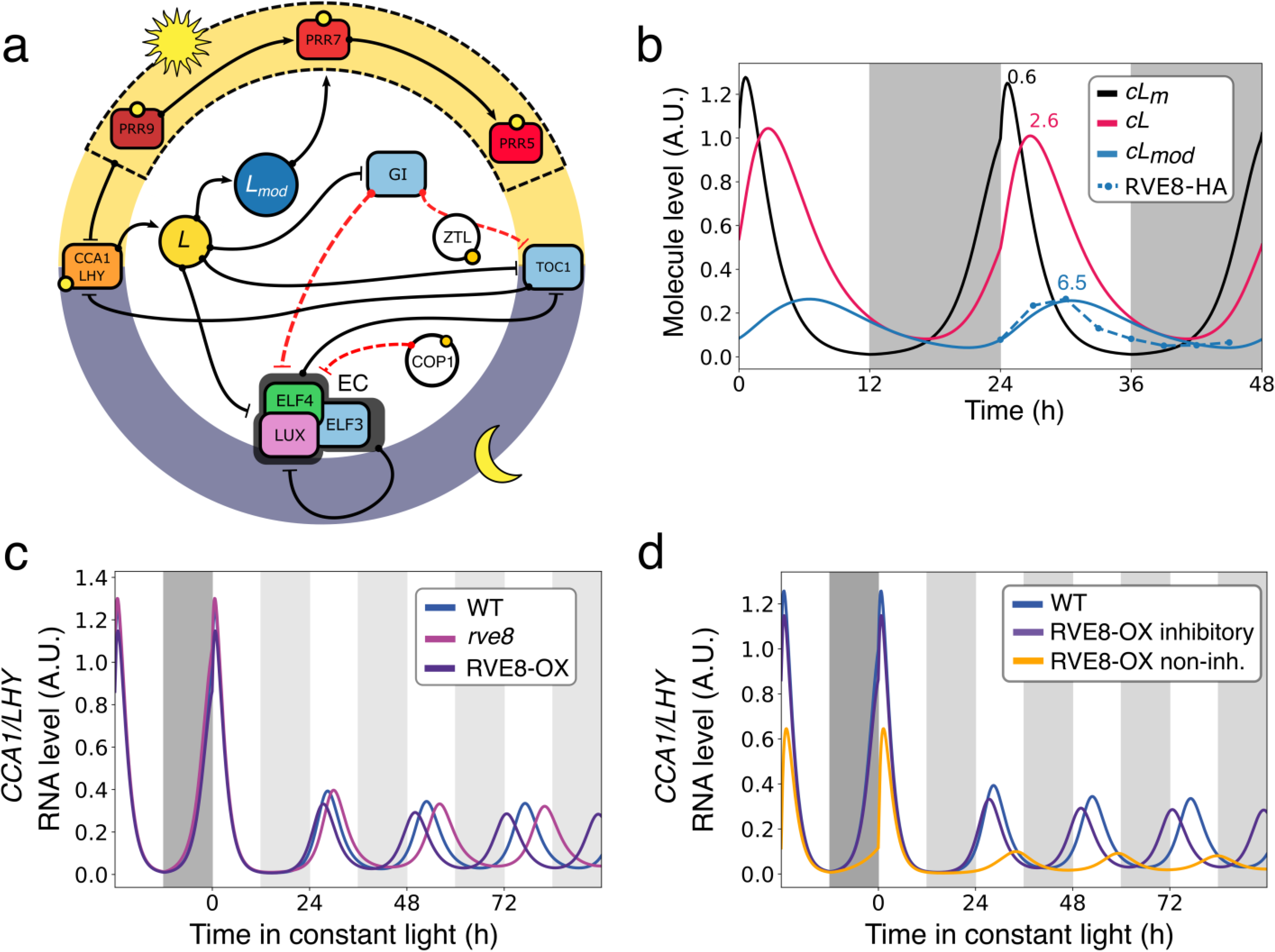
Transcriptional activation in P2011, compared to expression and function of *RVE8*. Simplified diagram locating *Lmod* in the P2011 model. Arrows, activation; blunt lines (T), repression. Solid edges, transcriptional regulation; dashed red lines, posttranslational regulation. Yellow circles, light regulation. The equivalent regulation of *CCA1/LHY* by each PRR protein is indicated by the dashed black box. b) P2011 simulation in L:D cycles (white:dark grey bars) showing the behaviour of CCA1/LHY variables *cL_m_* (mRNA), *cL* (protein), *cL_mod_* (modified, activating protein) and the similarity of *cLmod* to RVE8 protein data (RVE8-HA, (Hsu *et al*. 2013)). Peak expression times are indicated: 0.6 h *cL_m_*, 2.6 h *cL* and 6.5 h c*Lmod*. c) Simulated perturbations in P2011 that reproduce the rhythmic phenotypes of Col-0, *rve8* and *RVE8-OX* plants. The model was entrained for 10 days in L:D cycles (white:dark grey bars) before transfer to constant light (white:light grey) at time 0. d) Inhibiting *L* function is a necessary part of *Lmod* function. *RVE8-OX* was simulated by increasing the model parameter *p3* (as in c), purple line) including the inhibitory effect of *Lmod* formation that partly consumes *L*. In contrast, the *RVE8-OX* non-inhibitory simulation (non-inh., yellow line) generated the same level of *Lmod* without depleting *L*, and failed to match the *RVE8-OX* phenotype. Simulations were otherwise conducted as in c).

Starting at dawn, expression of the highly-homologous LATE ELONGATED HYPOCOTYL (LHY) and CIRCADIAN CLOCK ASSOCIATED 1 (CCA1) genes peak simultaneously. The myb-related LHY and CCA1 proteins in the plant repress the *PSEUDO RESPONSE REGULATOR* gene family (*PRR9, PRR7, PRR5* and *TOC1* <u>=</u> *PRR1*), and of GIGANTEA (GI), LUX-ARHYTHMO (LUX), EARLY FLOWERING 3 (ELF3) and EARLY FLOWERING 4 (ELF4) (Alabadí*et al.* 2001; Kamioka *et al*. 2016; Mizoguchi *et al*. 2002; Nagel *et al*. 2015; Nakamichi *et al*. 2010; Schaffer *et al*. 1998; Wang and Tobin 1998). The latter, ‘evening genes’ are repressed until LHY and CCA1 protein levels fall during the day, whereas the *PRR* genes are expressed in a sequence from the early morning through the day until after dusk (Fujiwara *et al*. 2008; Nakamichi *et al*. 2010), and repress the expression of *CCA1* and *LHY*. Matching the observed *PRR* dynamics in the precursor model of P2011 required a transcriptional activator function (Pokhilko *et al*. 2010). Some members of the *REVEILLE/LHY-CCA1-like* family of transcription factors, which are dawn-expressed genes homologous to *LHY* and *CCA1,* were later observed to function as transcriptional activators. Family members notably RVE8/LCL5 have been shown to interact with a family of transcriptional co-activators termed NIGHT LIGHT-INDUCIBLE AND CLOCK REGULATED GENES, the LNK1-LNK4 family (Rugnone *et al*. 2013), which were not represented in either model. The evening-expressed genes *LUX, ELF4* and *ELF3* together produce the hetero-trimeric protein Evening Complex (EC) (Nusinow *et al*. 2011), which binds to the promoters of *PRR9, TOC1/PRR1, LUX, ELF4* and *GI* and represses their expression, in turn allowing *LHY* and *CCA1* expression on the next cycle.

Both the P2011 and F2014 models were appropriate to consider for the incorporation of absolute mass units. Three factors led us to start from the P2011 model. First, F2014 is the most complex clock model (Supplementary Table 1), which makes model development more technically challenging. Second, F2014 includes genes such as *BOA/NOX* and *RVE8*, for which less data is available to constrain the models than for other genes, and in particular no data is available in absolute units. Third, the F2014 model fixes all translation rates to an arbitrary value, whereas our long-term aim requires the models to represent proteins in absolute units, with realistic translation rates. Our approach was therefore to re-factor the P2011 model to use absolute mass units. Our work towards that goal has generated two alternative models, U2019 and U2020. U2020 now includes key interactions of the F2014 model in a simpler context. We use multiple data sources from the literature to test aspects of the models that were not previously accessible, in particular the models’ inferred transcription rates.

## Methods

All experimental data have been published elsewhere. The key results for model fitting derived from the TiMet project funded under the EU Framework Programme 7, and are publicly available as described (Flis *et al*. 2015). Other data sources are cited in the text. All model analysis was performed using python 2.7, in the computational environment described below and available as described in the Reproducibility section.

### Model analysis and availability

The P2011 model was translated from MATLAB (Pokhilko *et al*. 2012) into the high-level Antimony language and then translated into SBML using the python package Tellurium (Choi *et al*. 2018). The model translations of P2011 are available in the public FAIRDOMHub repository at https://fairdomhub.org/assays/1219. The F2014 model was originally written in clocksim (http://cbbp.thep.lu.se/activities/clocksim/F2014-20140425.tar.gz) (Fogelmark and Troein 2014). As a contribution to the community, we also translated one of the parameterisations of the F2014 model into Antimony and SBML, available at https://fairdomhub.org/studies/508. The U2019 and U2020 models were developed and analysed using Tellurium. SBML files are available using the root path www.fairdomhub.org/models/. Each model can be accessed by appending the FAIRDOMHub ID listed below, for example U2019.1 can be accessed as www.fairdomhub.org/models/726, and similarly for U2019.2 (727), U2019.3 (728), U2020.1 (729), U2020.2 (730) and U2020.3 (731). Equations and parameter values were extracted for publication from these files using COPASI version 4.30.240 (please see Supplementary Information).

### Model fitting

To estimate parameter values by comparison to experimental data, SBML models were imported into the SloppyCell python package (Myers *et al*. 2007). Step functions in SloppyCell were included to simulate diurnal changes of light conditions. Models were entrained for 10 cycles of 24 hours with 12 h of light and 12 hours of darkness, and the match to RNA timeseries data was tested using the *χ^2^* statistic, as follows. The TiMet data set for RNA profiles in one Light:Dark cycle (LD) followed by constant light was compared to simulations for the same conditions on simulation days 11 to 13, with constant light from dawn on day 12. To test the simulated rhythmic periods, the period of the *cLm* variable (*CCA1/LHY* mRNA) was measured in simulated data starting from 276 h (12h after the start of constant light), except for the long-period *prr79* mutant which used data from 300 h, and the *cEC* variable was used for the *lhy/cca1* double mutant. The target period values for each genotype are described in Table 1.

**Table 1.** Period constraints enforced in model development. The target periods (Enforced) and the observed periods for the U2019.3 and U2020.3 models are listed, for the Col-0 wild type and the clock mutants tested by simulation. The set of target periods is retained from earlier model development (Flis *et al*. 2015).

| Genotype | Enforced (h) | U2019.3 (h) | U2020.3 (h) |
| --- | --- | --- | --- |
| Col-0 | 24.5 | 24.5 | 24.48 |
| *prr7/9* | 30 | 30.01 | 29.99 |
| *lhy/cca1* | 17 | 16.7 | 16.99 |
| *gi* | 22 | 22 | 22 |
| *toc1* | 21 | 20.99 | 20.98 |
| *ztl* | 27 | 27.01 | 27 |

To update the parameter values to better match experimental data, all the parameters in the model were allowed to vary except for the Hill coefficients and the set (*m11, m27, m31, m33, n14, n5, n6, p15, p6, p7*). This set contains parameters related to CONSTUTITIVE PHOTOMORPHOGENESIS 1 (COP1) which were inferred previously using data for ELONGATED HYOCOTYL 5 (HY5) and LONG HYPTOCOTYL IN FAR-RED (HFR1) proteins in separate experiments (Pokhilko *et al*. 2012), and parameters controlling the dynamics of the hypothetical protein P. Parameter sets were updated using the Levenberg-Marquardt method of SloppyCell. When fitting scaling factors, we used the sensitivity equation functionality built into SloppyCell, which updates efficiently by determining the direction of largest gradient change in the cost function landscape (Myers *et al*. 2007). For *ad hoc* constraints, such as period values and amplitude, SloppyCell provides prototype methods that were adapted to implement a gradient by finite difference approach. The TiMet timeseries data and period constraints were not sufficient to enforce oscillatory dynamics in constant light conditions, for the models described in this work, hence amplitude constraints were also included. The modified SloppyCell codebase was documented with git and is available from https://github.com/jurquiza/SloppyCell_Urquiza2019a. The modifications are highlighted in the code base with the string “Uriel”.

### Stepwise development of model U2020.1 from U2019.1

To replace the *PRR* activation cascade in model U2019.1 with a repression cascade, we introduced a new set of variables that represent the *PRR* genes, with the revised regulation. The repressive function of CCA1/LHY (*cL*) and its activating derivative (*cLmod*) were retained. The new equations did not initially feed back into the original equations, to avoid wholesale modification of the model’s behaviour. The parameters for these “dummy” equations were fitted to *PRR* profiles simulated from U2019.1 using the Levenberg-Marquardt algorithm of SloppyCell. The original *PRR* equations could then be replaced with the new equations, with only minor changes to model behaviour. The new model was then fitted to data simulated from U2019.1 for the entire set of variables, conditions and mutants present in the TiMet data set (leaving only parameters associated with COP1 and P fixed, as noted). The aim was to recover P2011 behaviour using the update network topology resulting in the model U2020.1

### Reproducibility

We developed a Docker image (uurquiza/urquiza2019a_tellurium_sloppycell:latest) to ensure the reproducibility of computational results. The image is publicly available from the Docker hub (https://hub.docker.com) and from https://fairdomhub.org/assays/1224. After installing the Docker software, the image can be downloaded by typing docker pull image_name. The image can also be built from source code available in the git repository https://github.com/jurquiza/Urquiza2020a. This repository contains instructions for installation and for running the Docker image, which compiles all the required tools for model fitting to oscillatory dynamics. The fitting is described in Jupyter notebooks in the same git repository.

## Results

Before re-factoring the P2011 model to use absolute units, two aspects of this model were reconsidered in the light of recent evidence and insights from the F2014 model.

### Transcriptional activation in the P2011 model and simulation of REVEILLE 8

We and others have focussed on the prevalence of negative regulation in the Arabidopsis clock circuit, and on transcriptional repression in particular. The P2011 model also includes transcriptional activation by the variable *Lmod* (denoted *cLmod* in written model equations, or cLm in P2011 in SBML format). *Lmod* activates the expression of the later-peaking *PRR* genes *PRR7* and *PRR5* (Figure 1a; *PRR5* is identified with the Night Inhibitor predicted by the P2011 model). *Lmod* was first proposed in the earlier P2010 model, where this hypothetical component was required to match the extended profile of *PRR* gene expression (Pokhilko *et al*. 2010). It is represented as a modified version of the model’s combined CCA1/LHY protein *L*. *Lmod* functions only as a transcriptional activator (Pokhilko *et al*. 2010), whereas the unmodified protein acts as the known repressor of evening genes, and to activate early *PRR* gene expression. CCA1 protein was known to be phosphorylated in a manner that altered DNA binding (Andronis *et al*. 2008), suggesting one potential biochemical mechanism for the modification. Slow conversion of *L* into *Lmod* leads to peak accumulation of *Lmod* at ZT 6.5 h (Figure 1b), providing a peak of activation at midday. The RNA variable that represents *CCA1* and *LHY* transcripts peaks just after dawn, in line with the experimental data. Therefore, the model includes a 6 h lag between the transcript peaking time and maximal accumulation of *Lmod*. This effective solution matched the (then) unknown regulation of the *PRR* genes and, given the lack of constraining evidence, it was presented as a parsimonious but phenomenological construct.

Interestingly, the REVEILLE 8 / LHY-CCA1-LIKE 5 (RVE8 / LCL5) gene product presents very similar behaviour (Rawat *et al*. 2011). The levels of *RVE8* transcript peak at dawn and RVE8 protein levels peak 6 h later (Figure 1b, RVE8-HA). In order to test whether *Lmod* in the model recapitulated the functions of *RVE8* in the plant, we sought to replicate two published perturbations of *RVE8*. The *rve8* mutant has a late rise of *CCA1* transcript under constant light conditions. Setting the transformation rate of *L* to *Lmod* (parameter *p3*) to zero in P2011 replicated the mutant phenotype (Figure 1d, *rve8*). This manipulation delays the repression of *CCA1* transcription and results in a later peak and long circadian period in constant light, also matching luciferase imaging data (Farinas and Mas 2011; Rawat *et al*. 2011). Experimental overexpression of *RVE8* (in *RVE8-OX* transgenic plants) results in period shortening (Farinas and Mas 2011; Rawat *et al*. 2011; Shalit-Kaneh *et al*. 2018), whereas simulating constant high levels of *Lmod* resulted in arrhythmia. Though *Lmod* resembled *RVE8* in the loss-of-function test, it did not recapitulate the simplest simulation of *RVE8* overexpression.

We therefore tested whether an alternative mechanism could explain the behaviour of *RVE8-OX,* reasoning that RVE8 might function as a required component of a hypothetical Noon Complex (NC) of proteins, which collectively form the transcriptional activator. If other components of the complex are also rhythmically regulated with peak abundance around ZT6, then *RVE8* overexpression might accelerate NC formation without altering its peak time. Increasing the transformation rate of *L* into *Lmod* (parameter *p3*) simulated this effect, and recapitulated the early rise of *CCA1* transcript that is observed in the *RVE8-OX* genotype, along with its short period (Figure 1c) (Rawat *et al*. 2011).

Experimental data from studies of the LNK protein family (Rugnone *et al*. 2013) is consistent with the existence of a rhythmic Noon Complex. The *LNK* gene family is expressed in the morning and mutants of *LNK* genes present slower clocks, similar to the *RVE* family (De Leone *et al*. 2019; Hsu *et al*. 2013). Interaction between RVE and LNK proteins is reported *in vivo* (Xie *et al*. 2014) but plants that constitutively express both *RVE8* and *LNK1* do not show continuous interaction between these proteins. Rather, co-immunoprecipitation timeseries show that their interaction is still phase-specific (Pérez-García *et al*. 2015) (supplementary data). This result suggests that a third, rhythmically expressed component is required for complex formation (see Discussion), in line with the proposed Noon Complex. *Lmod* was proposed on functional grounds and might represent a functional Noon Complex more closely than it represents the individual, molecular constituent RVE8.

However, increasing the *L* -> *Lmod* transformation rate by increasing the parameter *p3* both increases *Lmod* and decreases *L* concentrations. Therefore, we dissected this dual effect (Figure 1d). Simulating only the increase of *Lmod* accumulation without impacting *L* levels, by introducing a dummy parameter that scales *p3* only for the *Lmod* synthesis equation, did not recapitulate the RVE8-OX phenotype, indicating that the post-translational, inhibitory effect on CCA1/LHY function is required in the model. This might represent RVE8 binding competitively with CCA1/LHY to DNA or inhibiting co-repressor recruitment by CCA1/LHY in the plant. We note that genetic evidence from the quintuple mutant *lhy/cca1/rve468* has been taken to suggest antagonism between the function of these RVE proteins and CCA1/LHY (Shalit-Kaneh *et al*. 2018). Analysis of Protein Binding Microarray profiles (Franco-Zorrilla *et al*. 2014) showed a significant correlation between the DNA binding motifs of CCA1 and RVE1 (no data are available for RVE4, 6 or 8), whereas the binding site preferences of CCA1 and LUX differed more in the same comparison (Supplementary Figure 1) even though LUX shares the Evening Element target sequence. This result is not surprising, as proteins that share DNA binding domain sequences also tend to have similar binding motifs (O’Malley *et al*. 2016; Weirauch *et al*. 2014). Nonetheless, our analysis supports the notion that competition for DNA binding might mediate the observed genetic antagonism, and our simulations suggest that this or another inhibitory mechanism contributes to the phenotype of RVE8-OX plants. The existing behaviour of *Lmod* concisely represents these effects. There is insufficient justification to complicate the model by representing the Noon Complex explicitly, with a time-constrained interaction of RVEs, LNK1 and an unknown, third component(s) (see Discussion).

### Formation of the Evening Complex and its role in the *lhy;cca1* double mutant

Mapping this transcriptional activation function to *Lmod* in the P2011 model has further implications for modelling studies. First, it suggests a different way to simulate the *lhy;cca1* double mutant (Supplementary Figure 2). The short-period, damped rhythms observed in this mutant have been important in model development, because model circuits must retain at least one rhythmic feedback loop in the absence of *LHY* and *CCA1* function, in order to match this observed behaviour (James C. W. Locke *et al*. 2005). Even with an appropriate regulatory circuit, few sets of parameter values retain this behaviour, and simulations of the double mutant are arhythmic in several clock models (Flis *et al*. 2015; Fogelmark and Troein 2014; Zeilinger *et al*. 2006). However, previous studies simulated the double mutant by eliminating translation of the *cLm* RNA. This removes both the *L* repressor functions and also the activator *Lmod*. Below, we simulate the *lhy;cca1* double mutant by eliminating only the repressive connections from *L* to its target genes, leaving the activating function of *Lmod* intact.

A second update involves the key clock component that remains in the *lhy;cca1* double mutant, namely the heterotrimeric Evening Complex (EC) (Nusinow *et al*. 2011). In P2011, this is formed when ELF3 and ELF4 form a heterodimer, and interaction with LUX completes the EC. The complex acts as a repressor of *PRR9, TOC1, LUX and ELF4* (Chow *et al*. 2012; Ezer *et al*. 2017)(Figure 1a). Feedback of the EC on its constituent *LUX* and *ELF4* genes remains rhythmic in simulations of the *lhy;cca1* double mutant (Supplementary Figure 2). Protein-protein interaction processes typically operate at a much faster timescale (milliseconds to seconds) than the transcriptional dynamics for the clock (minutes to hours). The P2011 model used this separation of timescales to introduce quasi-steady state assumptions for several steps in EC formation, reducing the number of sub-complexes that were modelled explicitly and hence the number of components and equations in the model. We revisited these assumptions, which imply that the rates of the constituent processes are fast. In P2011, however, the equations representing the dimerization of ELF3:ELF4 suggest that the complex never dissociates, and a combination of parameter values results in very slow formation of the EC, leading to a peak of complex formation in the middle of the night. (Supplementary Fig. 2a). The slow formation of the EC contributes to rhythmicity in the *lhy;cca1* double mutant (Supplementary Fig. 2b). These slow dynamics can be maintained by representing the sub-complexes involved in EC formation explicitly, increasing the complexity of the model but avoiding the constraining assumptions. The baseline U2019 model for this study retained the arbitrary mass units as in P2011 but used the insights and updates outlined above, which should also be relevant to other clock models (Fogelmark and Troein 2014; Pokhilko *et al*. 2013).

### Development of the U2019.1 model

The updated model equations were initially fitted to simulation results from P2011 using the SloppyCell software (Myers *et al*. 2007), in order to recreate dynamics similar to P2011. The fitting process used simulated wild type plants (WT), and the mutants *lhy/cca1, prr7/9, toc1, gi* and *ztl*. The *ztl* mutant is important for constraining the decay rate of TOC1 protein, though it is absent from the TiMet RNA timeseries data used below. This first model takes the name U2019.1 (available from https://www.fairdomhub.org/models/726). Figure 2a outlines the nomenclature of subsequent model versions, while Figure 2b shows the contributing software and resources.

**Figure 2.**
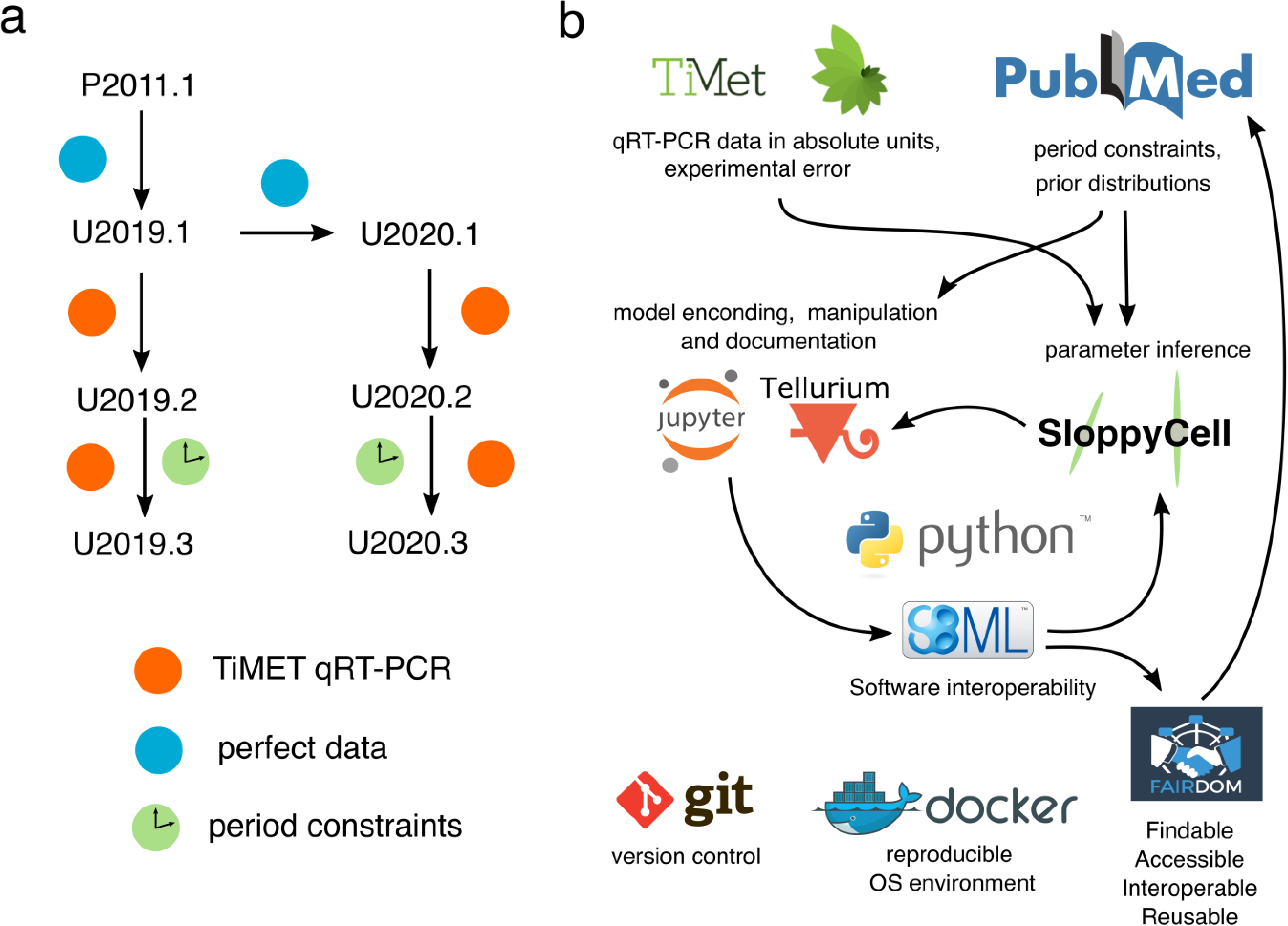
Model development with open and reproducible resources. a) Model nomenclature. Model architectures are denoted by author surname and year, with versions or parameter sets after a decimal point(s) (Flis *et al*. 2015). The published P2011.1 model simulated perfect data (blue). The U2019.1 model derived from P2011.1, without its quasi-steady state assumptions. U2020.1 has the *PRR* repression cascade (Figure 3 and main text). Parameters of both U2019.1 and U2020.1 were fitted to the perfect data of P2011.1. Models U2019.2 and U2020.2 were obtained by rescaling only transcription and translation rates such that modelled RNA levels matched TiMet RNA data (red). The U2019.3 and U2020.3 models resulted from a near-global re-optimisation to TiMet RNA timeseries, period and amplitude constraints (green). b) Reproducibility and wide reuse of models requires open data analysis tools. Public reference data of the TiMet project and other literature resources informed model development in open languages such as Python and Jupyter. The SBML model exchange format is supported by an ecosystem of open-source software for systems biology, such that models can be written in Antimony, explored in Tellurium, and fitted in SloppyCell. Models can then be transferred to Tellurium for further analysis. FAIRDOMHub and GitHub are crucial for accessibility and version control. We use Docker and DockerHub to disseminate our software environment reproducibly.

### Structural changes for *PRR* regulation (backward temporal inhibition) in U2020.1

In P2011 the wave of PRRs is a consequence of mutual activation, in order CCA1/LHY (*Lmod*) -> PRR9 -> PRR7 -> PRR5 (Figure 1a). Experimental evidence suggests repressive activity for the PRRs (Nakamichi *et al*. 2012, 2010; W. Huang *et al*. 2012; Gendron *et al*. 2012; Liu *et al*. 2013), suggesting a mechanism in which a wave of inhibition CCA1/LHY (Lmod) |-- PRR9 |-- PRR7 |-- PRR5 |-- TOC1 can produce similar peak times (Figure 3a). This mechanism was partially introduced in the P2012 model, where TOC1 inhibits *PRR9* transcription (Pokhilko *et al*. 2013). Other modelling efforts expanded this idea to further PRRs (Fogelmark and Troein 2014; Foo *et al*. 2016). Analysis of Foo et al. (2016) showed that repression results in cuspidate (sharp) peaks of gene expression, which could generate larger rhythmic amplitudes. High-amplitude regulation (100- to 1000-fold changes in RNA level) was a striking feature of the TiMet RNA timeseries, suggesting this mechanism could improve the model. Therefore, we updated the *PRR* activation chain to a repressive mechanism (Figure 3a), in a conservative, stepwise approach (see Methods), ending with a broad re-optimisation of parameters to fit data simulated from U2019.1. Figure 3b shows simulated *PRR7* RNA timeseries from an intermediate stage. Through this approach, the repression-wave model U2020.1 recovered behaviours similar to the activation-wave model U2019.1, illustrated by the simulated *PRR7* transcript (Figure 3c), which also shows the short period expected in simulations of the *toc1* mutant. The U2020 model thus represents an intermediate between P2011 and the greater complexity of F2014 (Fogelmark and Troein 2014). With the U2019 and U2020 model circuits in hand (Supplementary Figure 3), we next develop a method to use transcript data in absolute units to recalibrate the plant clock models.

**Figure 3.**
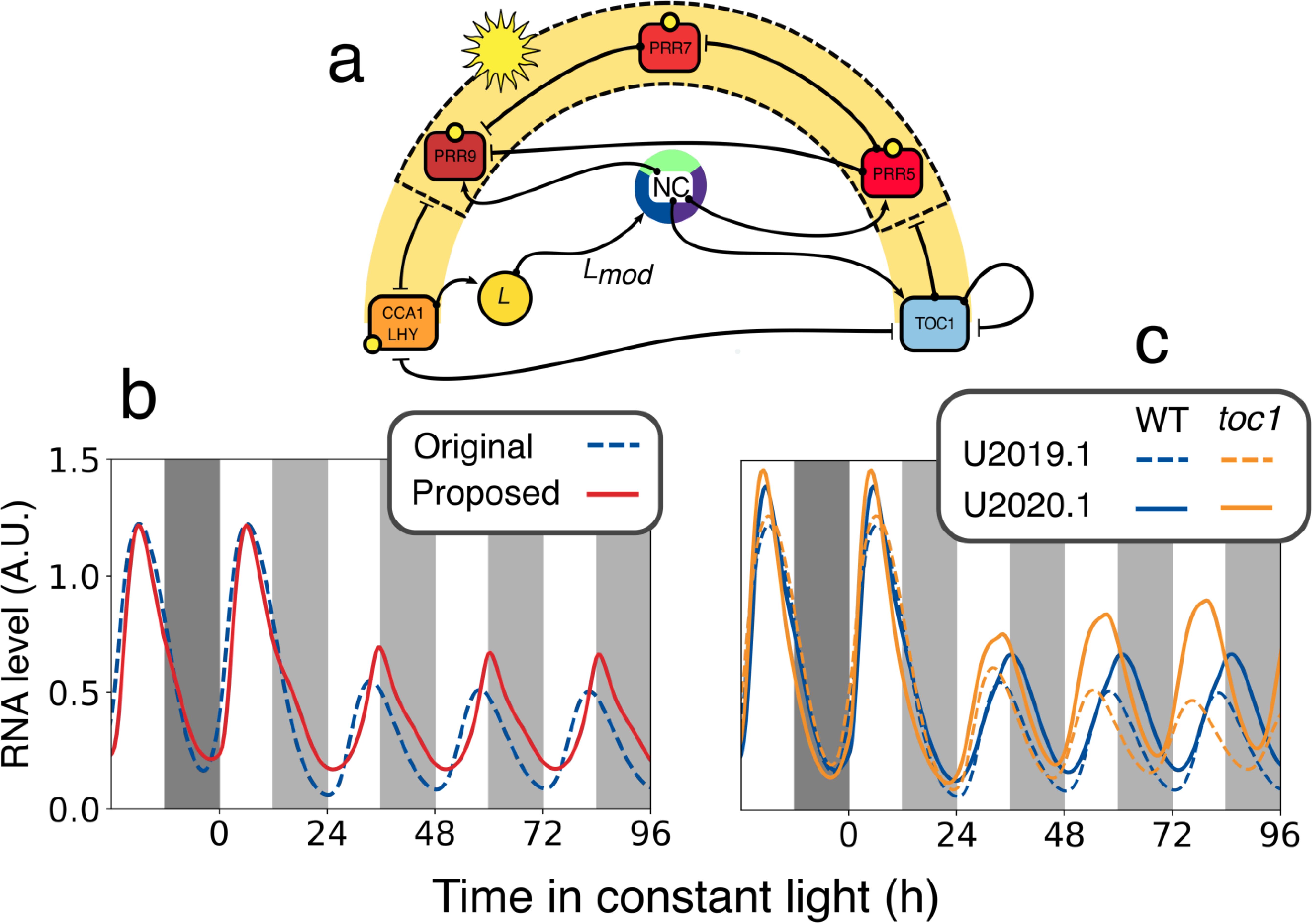
The refactored, repression cascade of *PRR*s in the U2019 and U2020 models. a) Simplified diagram of the *CCA1/LHY* and *PRR* gene families in the U2020 model, using the same conventions as Figure 1a. The Noon Complex (NC) proposed to function similarly to *Lmod* is shown as a pie, potentially formed from RVE, LNK and at least a third unknown component. b) Dynamics of *PRR7* transcripts in dummy variables controlled by U2019.1 under LD cycles then constant light; original, U2019.1 activation model, as in P2011 (dashed blue line); proposed, dummy variable for repression model (solid red line). c) Dynamics of *PRR7* transcript in U2019.1 (dashed lines) and U2020.1 (solid lines), in WT (blue) and *toc1* mutant (yellow). The derivation of U2020.1 is described in the main text. The models were entrained for 10 days in L:D cycles (white:dark grey bars) before transfer to constant light (white:light grey) at time 0.

### Data description

Flis et al (2015) performed absolute quantification of clock RNA transcripts, reporting data as average RNA copies/cell. The transcript collection consists of *LHY, CCA1, PRR9, PRR7, PRR5, TOC1, GI, LUX, ELF3, ELF4* and *GAPDH*. Timeseries studies that follow the emerging recommendations for quantification of these transcripts (Hughes *et al*. 2017) were performed in WT plants and four clock mutants (*lhy-21/cca1-11, prr9-11/prr7-11, toc1-101 and gi-201*). This provides a reference dataset for the transcriptional dynamics of plant clock models. Moreover, these data do not require internal normalising transcripts, reducing the risk of waveform distortion to the clock transcripts due to normalisation factors that might not remain constant over time.

### The experimental error in calibrated qRT-PCR data

Biochemically-realistic models in molecular biology generally present a large number of parameters. Experimental approaches for determining these parameter values directly can be extremely challenging. A model-based approach can be taken for estimating parameter values from time-series data of observed variables, such as the RNA timeseries used to develop the earlier clock models, discussed above. If the experimental error is well characterised, then a maximum likelihood approach can be taken. In particular, if the experimental error follows a normal distribution then the following likelihood function can be used,

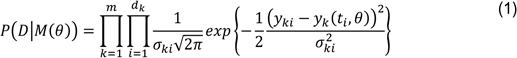

This measures the probability of data *D* being generated by a model *M* with a parameter vector *θ*. Experimental measurements are represented by *yki* with uncertainty *σki*. Model predictions for the measurements *yki* are given by *yk(ti,θ)*. The *k* index denotes state variables, while the *i* index represents time points. A maximum likelihood estimate for the model parameters can be found by minimising the difference between model predictions and data (Figure 4a), and this general approach has been widely used. However, the nature of the experimental uncertainty (‘error’) is important in the inference process and this has been relatively neglected. The nature of the errors depends on the methodology used for gathering the experimental data (Raue *et al*. 2013).

**Figure 4.**
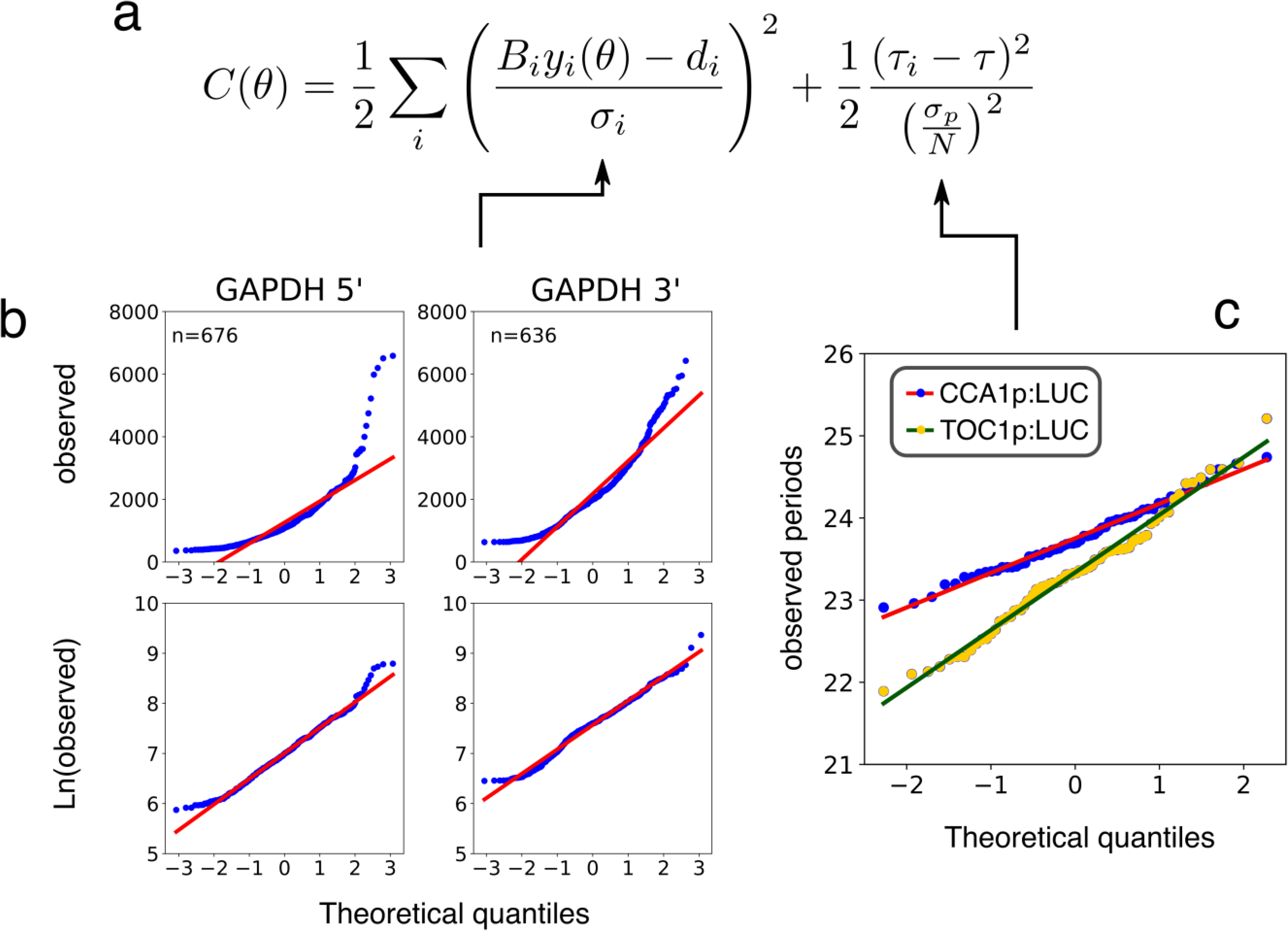
The cost function for model development uses experimental error estimates. a) The function used to calculate the cost *C(θ)*, for a parameter set *θ* compared to data from one genotype, sums costs for all datapoints *i*, comprising the departure of ln-transformed, simulated variable *y_i_(θ)* from the cognate, ln-transformed RNA timepoint *d_i_,* scaled by the experimental error estimate *σ* of that data point, and the departure of simulated period *tau(θ)_i_* from observed period *tau*, scaled by the experimental error estimate *σ_p_* of that period value and *N* the number of independent period estimates; here, *N* = 1. b) and c), the appropriate data transformation is inferred by the (mis)match between the distribution of a large set of (b) RNA or (c) period measurements and the expectation from the normal distribution (lines). b) quantile-quantile plots of GAPDH 5’ (left panels) or GAPDH 3’ RNA (right) data for untransformed (upper) and ln-transformed data (lower), with a better match in the latter. c) shows a good match to normality for untransformed period estimates from WT plants carrying a CCA1p:LUC (blue) or TOC1p:LUC (yellow) transgene.

Experimental error has been characterised in the context of the Arabidopsis clock, using RNA-seq data (Mombaerts *et al*. 2016). The TiMet data set used calibrated qRT-PCR data in units of [molecules] [cell]^-1^, which need not have the same error distribution. Though the method avoids normalisation to an internal reference RNA, nonetheless Flis et al (2015) also quantified the transcript level for a common reference gene Glyceraldehyde 3-phosphate dehydrogenase (GAPDH) using primers for both ends of the mRNA, in all the clock mutant backgrounds tested and across the 72 hours of sampling. These results for a nominally-constant RNA provide an opportunity to characterise the experimental error of this methodology.

We compared the raw qRT-PCR data of GAPDH to the expected theoretical quantiles for a normal distribution. We observed departure from normality at both tails of the distribution, for both the 5’ and 3’ UTR of GAPDH (Figure 4b). It has been reported that experimental error in the determination of molecular levels of components can be described by ln-normal models (Furusawa *et al*. 2005; Kreutz *et al*. 2007; Raue *et al*. 2013). Performing a ln-transformation on the data resulted in a better match to normality (Figure 4b). Therefore, data was ln-transformed for evaluating model costs. Furthermore, thanks to the large number of data points reported by Flis et al (n > 600), we were able to estimate the associated uncertainty (5’-UTR, σ = 0.95; 3’-UTR, σ = 0.34), which might be expected in other data similar to the TiMet study. The uncertainties derived from biological replicates at each time point are preferred (n=2 in the TiMet data). Where a replicate is missing, the inferred uncertainty (using the average of the 5’-UTR and 3’-UTR values) provided a reasonable proxy.

### Period constraints

Circadian period estimates are the second data type used for model fitting, and these are available for the WT and for several mutants. It has been generally assumed that the error in period estimates follows a normal distribution. To test this, we analysed publicly available period estimates for transgenic Arabidopsis plants that report that transcriptional activity of two clock promoters, CCA1p:LUC and TOC1p:LUC, tested in longitudinal imaging assays. The data were deposited in Biodare (https://www.biodare.ed.ac.uk/, experiment ID 12730563219125). Testing for departure from normality results in non-significant measures both for CCA1p:LUC (statistic=1.01, p-value=0.6) and for TOC1p:LUC (statistic=0.078, p-value=0.67), (Figure 4c) (R. D’Agostino and Pearson 1973; R. B. D’Agostino 1971). Therefore, the period data are well represented by a normal distribution, with σ = 0.52 for CCA1p:LUC and σ = 0.71 for TOC1p:LUC. Our period constraints derived from imaging studies using CCA1p:LUC, so the cognate standard deviation was incorporated into the likelihood function.

### Linking U2019.1 and U2020.1 to TiMet time-series and period constraints

With confidence in our normality assumptions, we next linked the models to the TiMet data set. First, we introduced mass scaling factors for each transcript variable. These result only in a change in the units of RNA variables, which is obtained by multiplying each transcription rate by its scaling factor and dividing the cognate translation rate by the same factor. We fitted the scaling factors of U2019.1 and U2020.1 to ln-transformed TiMet RNA timeseries data using SloppyCell. The resulting, rescaled models are named U2019.2 and U2020.2, and have units of [transcripts][cell]^-1^ for the transcript variables (Supplementary Figure 4). Up to this point, however, the dynamic behaviour of the models was constrained by the behaviour of the P2011 model, not by the experimental timeseries.

The rescaled models were then fitted to the TiMet RNA timeseries for WT, *lhy/cca1, prr7/9, toc1* and *gi* mutants, and to the corresponding period estimates of transgenic plants with these genotypes in constant light (Table 1). The period of the *ztl* mutant was again included to constrain the degradation rates of TOC1 protein. The resulting models are named U2019.3 and U2020.3, derived from U2019.2 and U2020.2 respectively. Figure 5 shows a sample of the fitted time-series, from simulations which correspond to Col-0 WT data for *LHY*, *PRR7, TOC1* and *LUX*. Table 1 shows that both models matched the target periods very closely. Furthermore, both models corrected the phase delay that the P2011 model showed upon transfer from Light:Dark (LD) cycles to constant light (compare Figure 5 to Figure 1d), when the model’s rapid degradation of PRR proteins in darkness is lost (Flis *et al*. 2015). The repression-based model U2020.3 shows a higher amplitude of RNA regulation, closer to the data for *LHY*, *PRR7* and *LUX* (see Discussion). Overall, we observe a slightly better fit to the data for U2019.3 (total cost 6.16×10^6^) compared to U2020.3 (total cost 1.02×10^7^), and U2019.3 has slightly fewer parameters (99 compared to 103, Supplementary Table 1). Updating the regulatory mechanisms did not guarantee an improved fit to the data.

**Figure 5.**
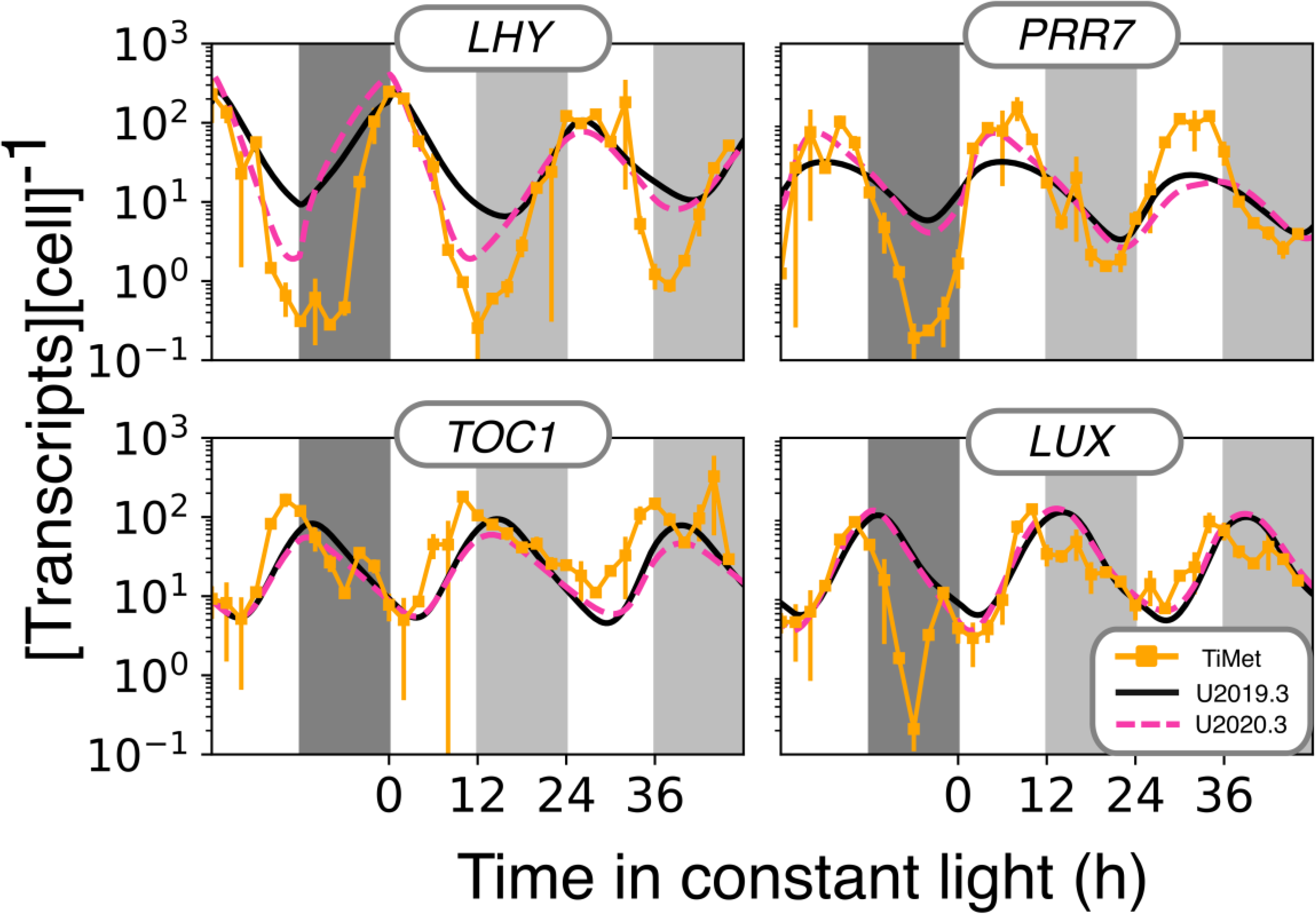
Behaviour of the U2019.3 and U2020.3 models, compared to TiMet RNA data for WT plants. RNA levels for four transcripts in Col-0 WT plants are shown, compared to simulation results from U2019.3 (solid black line) and U2020.3 (dashed red line). The higher amplitudes of *LHY* and *PRR7* data are better matched by U2020.3. The models were entrained for 10 days in L:D cycles (white:dark grey bars) before transfer to constant light (white:light grey) at time 0. Errors bars for experimental data are 1 S.D., n=2.

Table 2 shows the cost for each individual RNA timeseries in U2019.3 compared to U2020.3, highlighting which aspects of model behaviour were affected by the change in *PRR* gene regulation. The U2020.3 simulation of the Col-0 WT returned a better match (lower cost value) than U2019.3 for the genes *TOC1, PRR7, LHY, LUX* and *GI*. However, the absolute cost of *PRR9*, *LHY* and *GI* is higher for both models than for example *LUX* in U2020.3, where the simulated expression profile presents a notably good match to the high-amplitude regulation in the TiMet data (Figure 5).

**Table 2.**
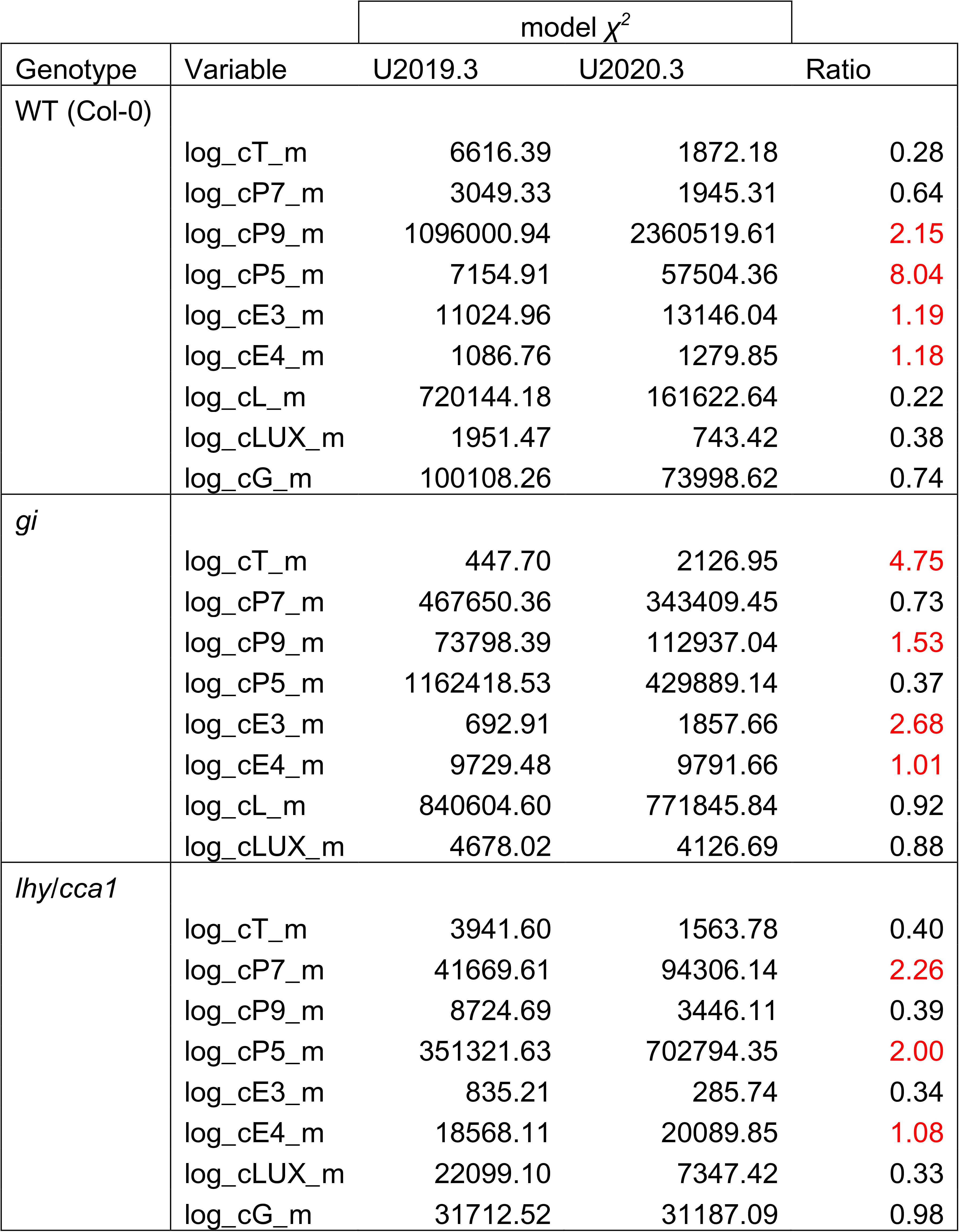
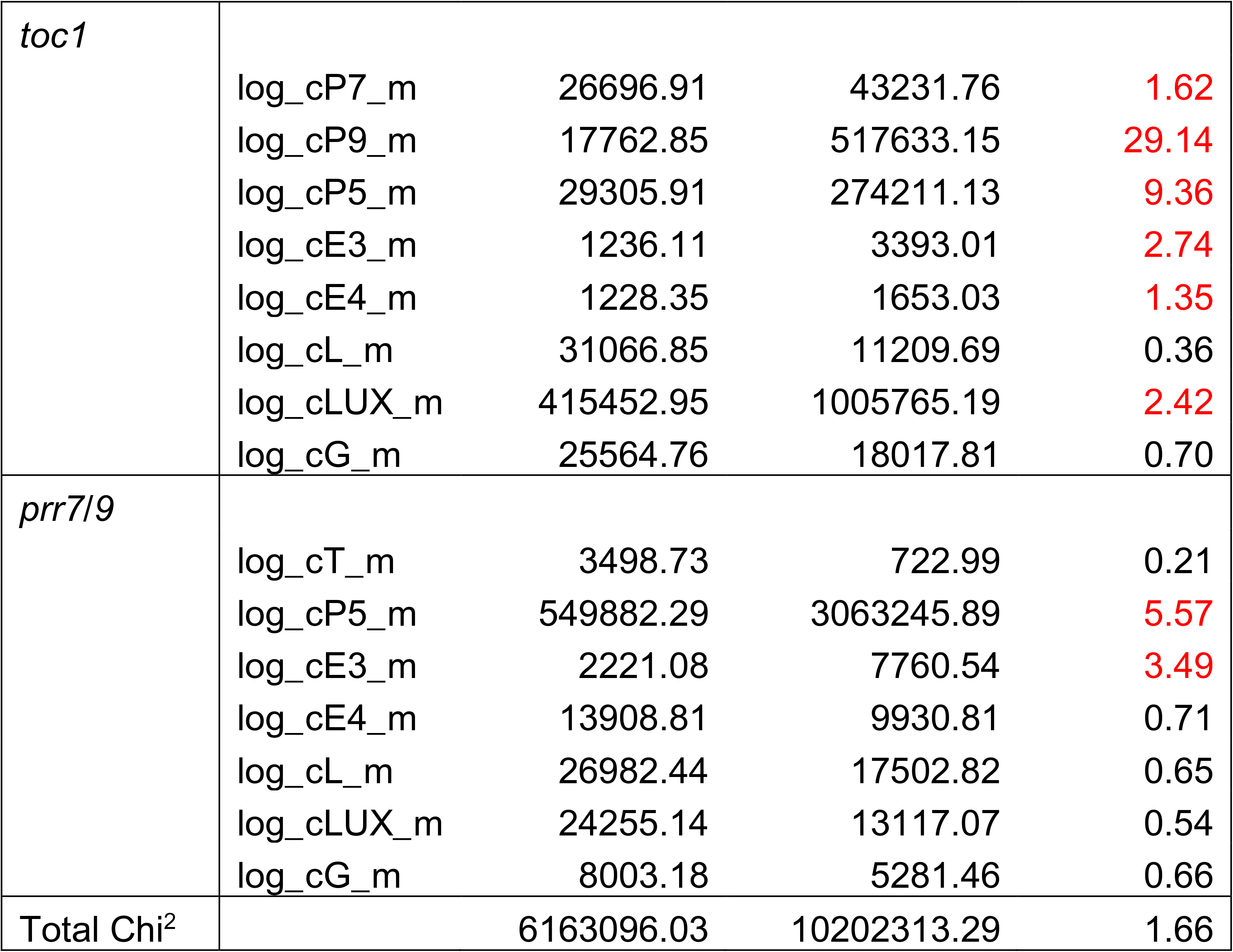
Cost function values of the U2019.3 and U2020.3 models compared to the RNA time series. Higher *χ^2^* values reflect a greater departure of each simulated RNA variable from the cognate TiMet RNA data, under one LD cycle followed by LL. A higher Ratio between these values reflects a greater departure of the U2020.3 simulation from the data, compared to U2019.3. Variable names are listed as the SBML variable identifiers; the cognate RNA data are as follows: cT_m, *TOC1*; cP7_m, *PRR7*; cP9_m, *PRR9*; cP5_m, *PRR5*; cE3_m, *ELF3*; cE4_m, *ELF4*; cL_m, *LHY*; cLUX_m; cG_m, *GI*. Some RNA species are absent in the mutants. The ‘log’ prefix denotes the ln-transformation of both data and simulated values.

Compared to the data for clock mutants, simulation of the *lhy/cca1* double mutant improved the fit for several variables in U2020.3, with moderately increased costs for *PRR7* and *PRR5*. The *gi* mutant data was also better matched by U2020.3, with a particular improvement in the simulation of *PRR5*. U2019.3 matched better than U2020.3 to the data for the *prr7/9* mutant, where the match of U2020.3 to data for *PRR5* expression had the greatest cost of all the timeseries. At the other end of the *PRR* cascade, the *toc1* mutant data was also better matched by the U2019.3 model, with *PRR9, PRR5* and *LUX* contributing to higher costs for U2020.3. Across all the genotypes, the regulation of these three transcripts would merit further attention, based on the fitting data from U2020.3.

### A genome-wide transcription rate distribution in absolute units

The recalibrated models now include transcription rates in absolute units that are compatible with the TiMet data and period constraints. The values of these model parameters (or strictly, the rates that are realised in simulation) can be taken as predictions of the functional rates *in vivo*. To provide a first-order test of these predictions, we constructed an empirical probability distribution of Arabidopsis transcription rates in units of [transcripts][cell]^-1^[h]^-1^, matching the model transcription rate parameters. No such data existed but two published data sets could be linked to derive the distribution, using simple assumptions.

The first data set consists of microarray transcriptomic measurements of synthesis and decay of RNAs in Arabidopsis, obtained in the BBSRC ROBuST project using the labelling kinetics of RNA with the ribonucleotide analogue 4SU (Sidaway-Lee *et al*. 2014), hereafter termed the Sidaway-Lee data (Figure 6b). This data includes only 7,291 genes which represents 35% of the 20,568 genes in the Arabidopsis genome (BNID 105446 https://bionumbers.hms.harvard.edu). The distribution has units of [microarray units] [cell]^-1^[h]^-1^. In order to transform the units of the distribution into [transcripts][cell]^-1^ [h]^-1^, we calibrated the Sidaway-Lee data using the second published data. The ‘Piques data’ comprises measurements at dawn and dusk of 96 transcripts coding for enzymes of Arabidopsis central metabolism (Piques *et al*. 2009), from the same, calibrated qRT-PCR method as (Flis *et al*. 2015). We selected a subset of nine transcripts that were measured in both data sets and that changed less than 0.2-fold in level between dawn and dusk in the Piques data (Figure 6a). Assuming that no dynamic regulation of their transcription rates was taking place in either study, the data should be equivalent.

**Figure 6.**
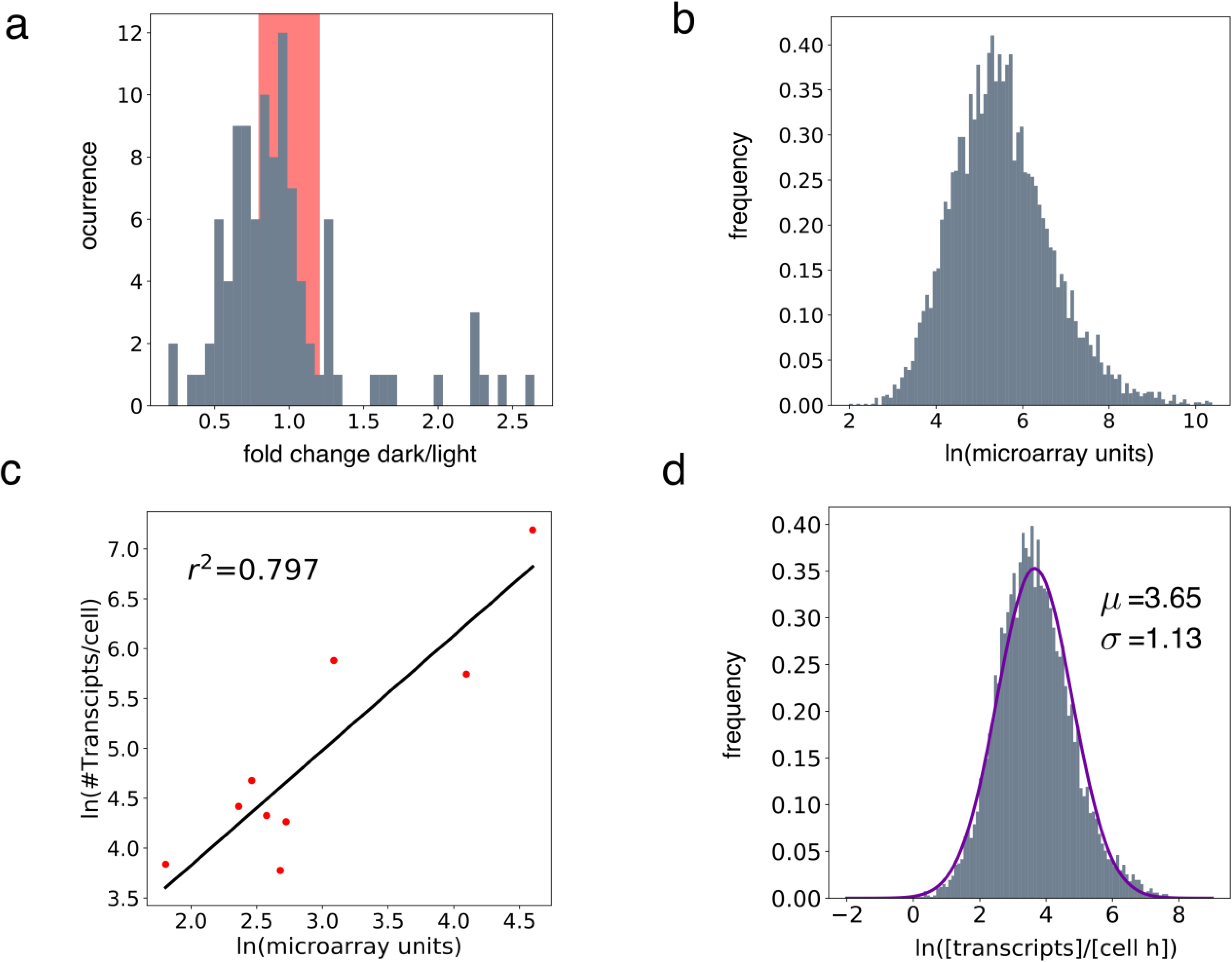
Recalibration of the Sidaway-Lee data into units of transcripts cell^-1^ h^-1^. a) Distribution of expression fold-change values between the two timepoints, for the 96 Arabidopsis transcripts in the Piques data set. The red band highlights the interval of selected transcripts, 1± 0.2. b) Steady state levels of transcripts at 17°C in the Sidaway-Lee data. c) Linear regression for the ln-transformed levels of selected transcripts from the Piques data against their levels in the Sidaway-Lee data. d) Sidaway-Lee’s distribution of transcription rates, transformed using the regression into units of ln[transcripts] [cell] ^-1^ [h]^-1^, compared to a normal distribution (solid line) with maximum likelihood estimation for the mean (μ=3.65) and standard deviation (σ=1.13) of the transcription rates.

The recalibration starts from a simple, dynamic model for the production of each transcript,

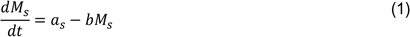

where *Ms* stands for the transcript level, *as* stands for the transcription rate and *b* for the transcript decay rate, and the subindex *s* refers to the Sidaway-Lee set. The units in this equation are [microarray units] [h]^-1^. We can model a quasi-steady state by setting the differential equation to zero and solving for *as*,

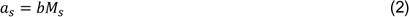

For the Piques data set the same model can be postulated, using parameter subindex *p*, but the transcription rates *ap* have units of [transcripts][cell]^-1^[h]^-1^. The decay constant *b* of each transcript should be equivalent in the two data sets, with units [h]^-1^, so we omit the subindices. The link between the two data sets can be achieved by performing a linear regression of Sidaway-Lee vs Piques data sets. The resulting regression model is substituted in the Piques quasi steady state equation as follows,

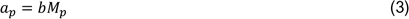

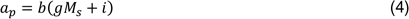

The gradient *g* has units of [transcripts][microarray unit]^-1^. We can now compare Sidaway-Lee data in [microarray units] [h]^-1^ on the right-hand side of the equation with Piques data in [transcripts][cell]^- 1^[h]^-1^ on the left-hand side. Figure 6c shows a highly significant correlation between the data sets (*r^2^* ∼ 0.8), and Figure 6d applies the regression to rescale the Sidaway-Lee distribution, yielding an average transcription rate of 38.4 in units of [transcripts][cell]^-1^[h]^-1^, with a one standard deviation span from 12.42-119.7 [transcripts][cell]^-1^[h]^-1^. This distribution should significantly aid the assessment of transcription rates in plant gene network models. We refer to it below as the Plant Empirical Transcription Rate distribution (PETR, pronounced Peter).

The linear regression approach described above can be tested using data sets that we expect to deviate more from our assumptions, where the correlation *r^2^* should drop. Fortunately, Sidaway-Lee et al described microarray studies for plants grown at two different temperatures 17 °C and 27 °C. The 17 °C data is used above as those conditions are closer to the conditions used by Piques et al. Encouragingly, the 27 °C data for the same genes returns a lower correlation coefficient (*r^2^* = 0.4) (Supplementary Figure 5, compare to Figure 6c). This result gives some support for the regression model used to develop the PETR distribution.

### Challenging predicted transcription rates with the PETR distribution

We tested the fitted and scaled clock models U2019.3 and U2020.3 by comparing the maximum transcription rates reached in simulations to the PETR distribution (Figure 7a, 7b). In both models, the transcription rates fall entirely within the distribution, with few in the tails. *cGm* transcription has a relatively high maximum (fast rate) compared to the other clock genes, for both models. This could reflect the requirement for strong, acute light activation of *GI* at dawn. In the case of U2019.3, CCA1/LHY presents a similarly fast maximum transcription rate, which is lower in U2020.3. For U2020.3, *PRR7* presents a rate among the highest estimates in the PETR distribution. These results indicate that the model circuits developed without the constraints of absolute mass units nonetheless predicted biochemically feasible transcription rates, potentially due to constraints on the cognate RNA degradation rates.

**Figure 7.**
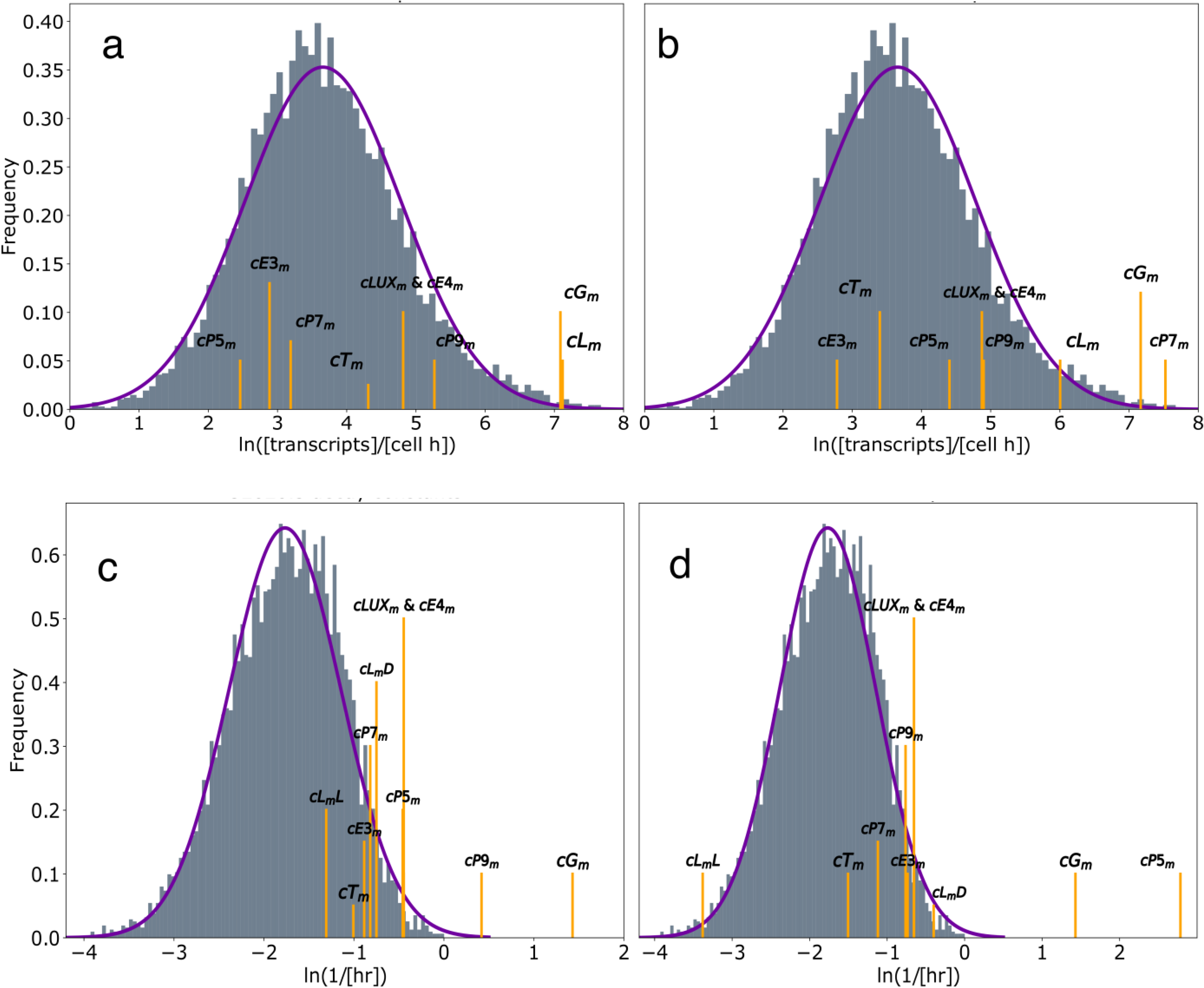
Parameters of RNA metabolism inferred in the models, compared to the observed distributions. a), b) Maximal transcriptional rate parameters relative to the PETR distribution and c), d) RNA degradation rate constants relative to the rates measured by (Sidaway-Lee *et al*. 2014), for the U2019.3 model (left panels) and U2020.3 model (right panels). *cL_m_L* and *cL_m_D* indicate decay rates for *cL_m_* in light and in darkness respectively. Yellow bars indicate the parameter value for each transcript variable. Solid lines show fitted normal distributions with unitless parameters μ=3.65, σ=1.13 (a, b) and μ=-1.76, σ=0.62 (c, d).

### Testing the RNA degradation rates of models against the Sidaway-Lee distribution

The 4SU-labelling approach also estimated RNA degradation rates (Sidaway-Lee *et al*. 2014), so the distribution of these observed rates can likewise test the inferred decay rate constants in the U2019.3 and U2020.3 models (Figure 7c, 7d). Again, most decay constants fall within the realistic range for Arabidopsis. The clock-associated genes tend to have transcripts with a short half-life, consistent with their dynamic regulation. The *LHY* RNA degradation rate in the light (*cLmL*) in U2020.3 is at the lower end of the distribution, whereas the rate in darkness (*cLmD*) is at the higher end. In the case of U2019.3, the *PRR9* RNA degradation rate is higher than any observed value. The *GI* RNA degradation rate in both U2019.3 and U2020.3 models also falls clearly outside the distribution and is likely to be unrealistically high. The *PRR5* RNA in the U2020.3 model presents the most extreme value of all the transcript degradation rates, and is not biochemically realistic. We offer further interpretation of these results in the Discussion.

## Discussion

Quantitative modelling of the Arabidopsis circadian clock mechanism has progressed significantly over more than 15 years (Bujdoso and Davis 2013). The complexity of the models has increased along with the inclusion of experimentally-documented variables (Supplementary Table 1). Naturally, most of the work has concentrated on simulating the timing of molecular events. The models have been challenged most stringently using timeseries data from mutant plants that alter rhythmic period and/or the waveform of clock components. Few of the inferred biochemical parameter values in the models could be tested against directly measured values, because neither the model parameters nor most of the available data were calibrated to absolute mass units that would allow direct comparisons. This work offers that calibration, for RNA levels and transcription rates.

### Interpretation of clock models

Some of the revisions of the P2011 model equations in the new U2019 and U2020 models were required to avoid earlier assumptions in the formation of the Evening Complex and had little effect on model dynamics, but unfortunately increased model complexity from 28 to 31 variables. In contrast, recognising the role of transcriptional activation has altered how we interpret and use the models, without increasing their complexity. In the P2011 model, the activation required for clock gene expression during the day was captured by a combination of CCA1/LHY protein and a modified version termed *Lmod*, which had to form slowly in the model to activate *PRR* gene expression later in the day. In particular, *Lmod* regulates *PRR5* in the models (Pokhilko *et al*. 2012). Given the later discovery and characterisation of the RVE and LNK families of transcription factors, we note the similarity between *Lmod* and RVE8 */* LCL5, particularly with respect to an enigmatic delay in observed RVE8 protein accumulation that matches *Lmod*. *PRR5* is also a key, direct target of RVE8 (Farinas and Mas 2011; Hsu *et al*. 2013; Rawat *et al*. 2011; Shalit-Kaneh *et al*. 2018; Xie *et al*. 2014). The activating function in our models was little discussed (Somers 2012), in part because it pre-dated the most relevant data.

Both the U2019 and U2020 models matched well to the target period values and RNA timeseries in wild-type and mutant plants (Tables 1 and 2), though the models differ in the mechanisms of *PRR* gene regulation (Figures 1a and 2a). The U2020 model had a slightly higher fitting cost and larger parameter number, so U2019 would be preferred *a priori*. However, the regulatory interactions in U2020, derived from the F2014 model, are better evidenced. As the number of plant clock models grows, it will be increasingly important to compare and document their performance in detail. If this is a collaborative effort between the model developers, it may also reveal implicit/tacit knowledge (Davies 2001) that is codified in each model, and which is then applied to simulate specific biological scenarios, exemplified by our reinterpretation of the *lhy/cca1* double mutant.

Identifying *Lmod* as a model of the observed, daytime transcriptional activator has a technical consequence, that simulations of the *lhy/cca1* double mutant in several models (P2010, P2011, P2012, F2016) should not remove all CCA1/LHY protein but instead should selectively remove its repressive effects. Simulating the damped, short-period rhythms in this mutant has been technically challenging (Zeilinger *et al*. 2006). The model retains very few feedback loops in this case, so its behaviour is highly sensitive to some of the parameters describing their biochemistry. The challenge remains when activating and repressing functions are represented as separate genes (Fogelmark and Troein 2014) or as effects on separate targets (this work; Flis et al., 2015). High parameter sensitivity is a marker for caution in modelling (Morohashi *et al*. 2002). The *lhy/cca1* double mutant simulations probably highlight the need for future work, which might have implications for our current view of the clock mechanism beyond this special case.

The biochemistry of transcriptional regulation contributes to the modelling challenges. Simulation of the clock dynamics in *RVE8-OX* plants encouraged us to propose the presence of a Noon Complex, formed by RVE8 and at least one other rhythmic partner, which might correspond more closely to the function of *Lmod* in the models. The LNK proteins are possible partners that interact with RVE proteins (Xie *et al*. 2014). However, in plant lines that constitutively express both RVE8 and LNK1, their interaction is still restricted to around ZT7 (Pérez-García *et al*. 2015), consistent with the requirement for at least one further, rhythmic partner. Affinity purification and mass spectrometry (AP-MS) could help in elucidating the cryptic component(s) required for NC formation and functionality, as applied to the Evening Complex (H. Huang *et al*. 2016) or in timeseries of interactions with the GI protein (Krahmer *et al*. 2019). Adding more of these processes explicitly will not only further complicate the models but also increase the uncertainty of their outputs, unless the new components are constrained by significant, additional data.

### PETR, a constraining distribution of transcription capacity for plant systems biology

By recalibrating our models into absolute units of transcripts per cell using the TiMet qRT-PCR timeseries, we made it possible to test the modelled transcription rates against biochemical data, if such data existed. Though we previously collated distributions of the binding affinities measured for plant transcription factors to DNA and for protein-protein interactions in Arabidopsis (Millar *et al*. 2019), we had relied on data from yeast to illustrate the possible transcription rates in plant cells. Integrating two apparently different data sets here resulted in the PETR distribution (Figure 6d), which estimates a mean transcription rate of 38.4 [transcripts][cell]^-1^[h]^-1^, with one standard deviation spanning 12.4-119.7 [transcripts][cell]^-1^[h]^-1^. Yeast for comparison has rates of 2 - 30 [transcripts][h]^-1^ (Pelechano *et al*. 2010). The higher mean in plants likely relates in part to the difference in size between yeast (43 µm^3^) and Arabidopsis cells (73,000 µm^3^) (Jorgensen *et al*. 2002; Wuyts *et al*. 2010). If the parameter values of other models are refactored into absolute units, the PETR distribution can be used to assess the resulting transcription rates *post hoc*. More likely, extensions to Bayesian inference, previously applied to the P2011 model (Higham and Husmeier 2013), might use the PETR distribution as a prior to constrain parameters to plausible values during model development. Directly measuring biochemical parameter values, including the absolute transcription rates of the clock genes, is still an important step for future model development.

The PETR distribution was derived as a log-normal distribution, following our assessment of the distribution of GAPDH qRT-PCR results in the TiMet data (Figure 4). Log-normal distributions are pervasive in biology, for example in western blot data (Kreutz *et al*. 2007). The statistical assumptions in parameter inference methods require a good description of the experimental noise, but it can be unclear what error distribution is appropriate, for example if time series have had to be extracted from data displays in the literature (Fogelmark and Troein 2014). This underlines again the importance of routinely sharing the numerical values of biological data (Milo *et al*. 2010).

### The implications of inferred clock gene transcription rates, and future model development

The key concerns in the parameter values governing the model’s RNA metabolism arose from the high RNA degradation rates predicted for some clock transcripts (Figure 7c, 7d). These reflect the timeseries data that the model fitting seeks to match, within the constraints of the model equations. Measured *GI* mRNA levels fall over 1100-fold in the 12 hours after dusk in the TiMet data, for example (Flis *et al*. 2015). The *GI* mRNA in the model has a constant half-life, which must be short enough to match these dramatic decreases in mRNA level. However, *GI* mRNA also accumulated 32-fold in the 2 hours after dawn (Flis *et al*. 2015), and even more rapid increases have been observed (Locke *et al*. 2005). Rapid accumulation of this very unstable transcript in turn requires a high, estimated *GI* transcription rate in the models (Figure 7a, 7b). For transcripts of *CCA1/LHY*, in contrast, the half-life is not constant as the model includes their observed, light-regulated degradation rate (Yakir *et al*. 2007). De-stabilisation of this mRNA in the dark helps to reach lower, trough levels in our model simulations (Figure 5) than in P2011 and earlier models (Figure 1b), more closely matching the data. Comparing our two models, the *CCA1/LHY* transcription rate is unusually high in the U2019.3 model but less so in U2020.3. One reason for this is likely that the difference between light and dark mRNA degradation rates is greater for the cognate RNA in U2020.3, in other words, the RNA is much less rapidly degraded in the light compared to darkness, lowering the demand on new transcription to match the observed RNA levels (Figure 7). Unusually high estimates of RNA metabolism parameters for *PRR* transcripts such as *PRR5* might also suggest that these RNAs are post-transcriptionally regulated *in vivo*, as is common in other clock systems (Martelot *et al*. 2012).

*PRR5* RNA expression not only has a high amplitude but also a cuspidate (peaked) waveform. Both are indicative of highly non-linear regulation, which might involve both regulated RNA stability and high-amplitude transcriptional regulation. The earliest modelling of circadian clocks used large Hill coefficients, which were later considered unrealistic, to simulate the cooperativity of transcriptional repressors. Our models conservatively limit the non-linearity of transcription factor binding, using Hill coefficients fixed at 2, which models the presence of a functional dimer. However, competition between alternative binding partners can also generate larger apparent Hill coefficients (Buchler and Louis 2008; Kim and Forger 2012; Lee and Maheshri 2012). Understanding such competition, for example among the RVEs and other transcription factors that bind the Evening Elements, will require the absolute quantitation of the copy number of regulatory proteins per cell, the number of genomic binding locations and the transcription factor affinity for those locations. Protein quantification is therefore another critical step for plant clock models but specific antibodies are available for only a few of the relevant proteins. Fusion proteins tagged using either firefly luciferase or NanoLUC offer a convenient alternative approach to protein quantification (Urquiza-García and Millar 2019).

Future digital organism models will require both simplified but still experimentally-based but models that facilitate analysis, and also biochemically-realistic models that link to genome sequences. These, detailed models should help to mobilise contributions from biochemistry, structural and chemical biology, which have influenced work on mammalian and cyanobacterial clocks far more than plant chronobiology. The detailed models will also assist the model-driven design of plant regulation, using precision genetic technologies such as Prime Editing (Lin *et al*. 2020).

## Acknowledgements

We gratefully acknowledge Anna Flis and Mark Stitt (Max Planck Institute for Molecular Plant Physiology, Potsdam-Golm, Germany) for providing raw qRT-PCR data ahead of publication, and Alastair Hume (EPCC, University of Edinburgh), Ryan Gutenkunst (University of Arizona, USA) and James P. Sethna (Cornell University, USA) for providing software support regarding *ad hoc* model constraints and the parallel processing capability in SloppyCell.

## Contribution by the authors

A.J.M. and U.U-G conceived the study and analysed modelling results. U.U-G. performed the model development, derived the PETR distribution, ensured software reproducibility and drafted the manuscript. U.U-G. and A.J.M equally contributed to the final manuscript.

The authors declare no conflicts of interest.

---

**Supplementary Table 1.** Models used or discussed in this work, including the numbers of parameters, variables and edges.

| Model | Derived from | # parameters | # edges | # state variables | used |
| --- | --- | --- | --- | --- | --- |
| P2011 | P2010 | 107 | 156 | 28 | yes |
| F2014 | P2011 | 119 | 182 | 35 | no |
| C2016 | P2011, F2014 | 43 + 9* | 37 | 9 | no |
| F2016 | P2011 | 81 | 102 | 24 | no |
| U2019.1 | P2011 | 99 | 156 | 31 | yes |
| U2019.2 | P2011 | 99 | 156 | 31 | yes |
| U2019.3 | P2011 | 99 | 156 | 31 | yes |
| U2020.1 | P2011, F2014 | 103 | 163 | 31 | yes |
| U2020.2 | P2011, F2014 | 103 | 163 | 31 | yes |
| U2020.3 | P2011, F2014 | 103 | 163 | 31 | yes |

\*Initial conditions are not usually considered as parameters, as they are irrelevant once the system has reached a stable limit cycle. The C2016 model has been shown to develop chaotic dynamics and quasi periodicity, in contrast, so initial conditions should be taken into consideration, increasing the effective parameter number.

**Supplementary Table 2.**
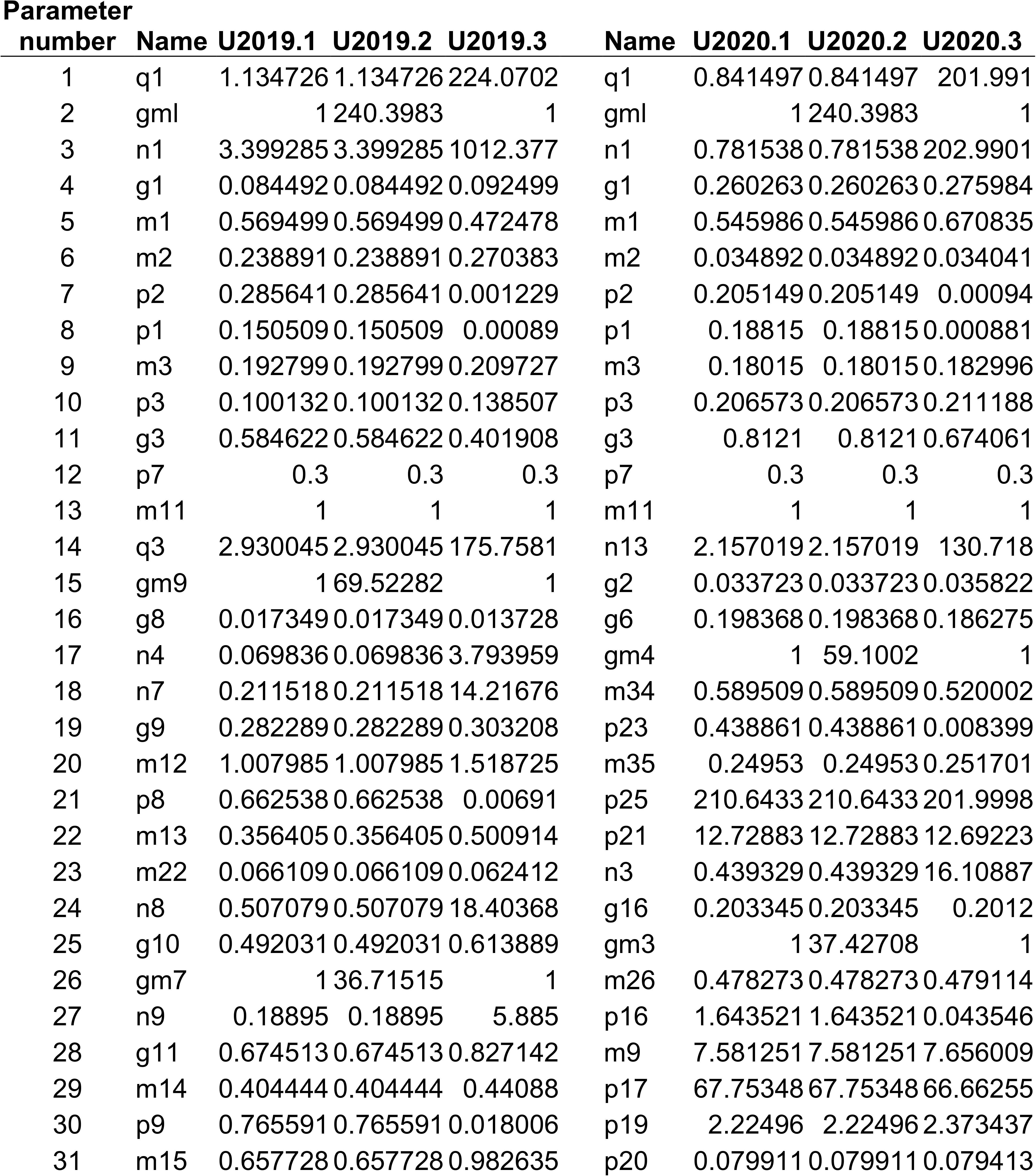
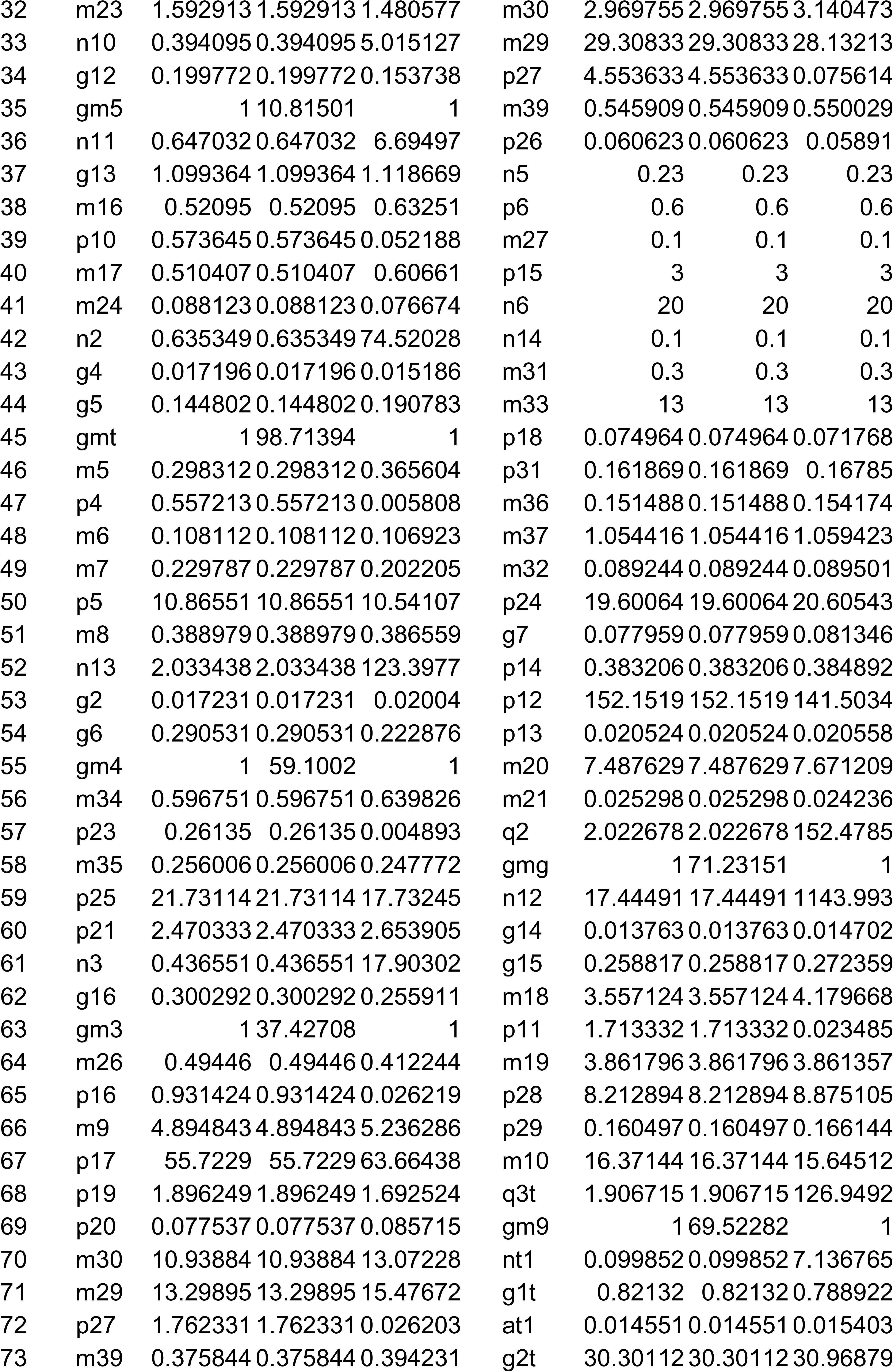
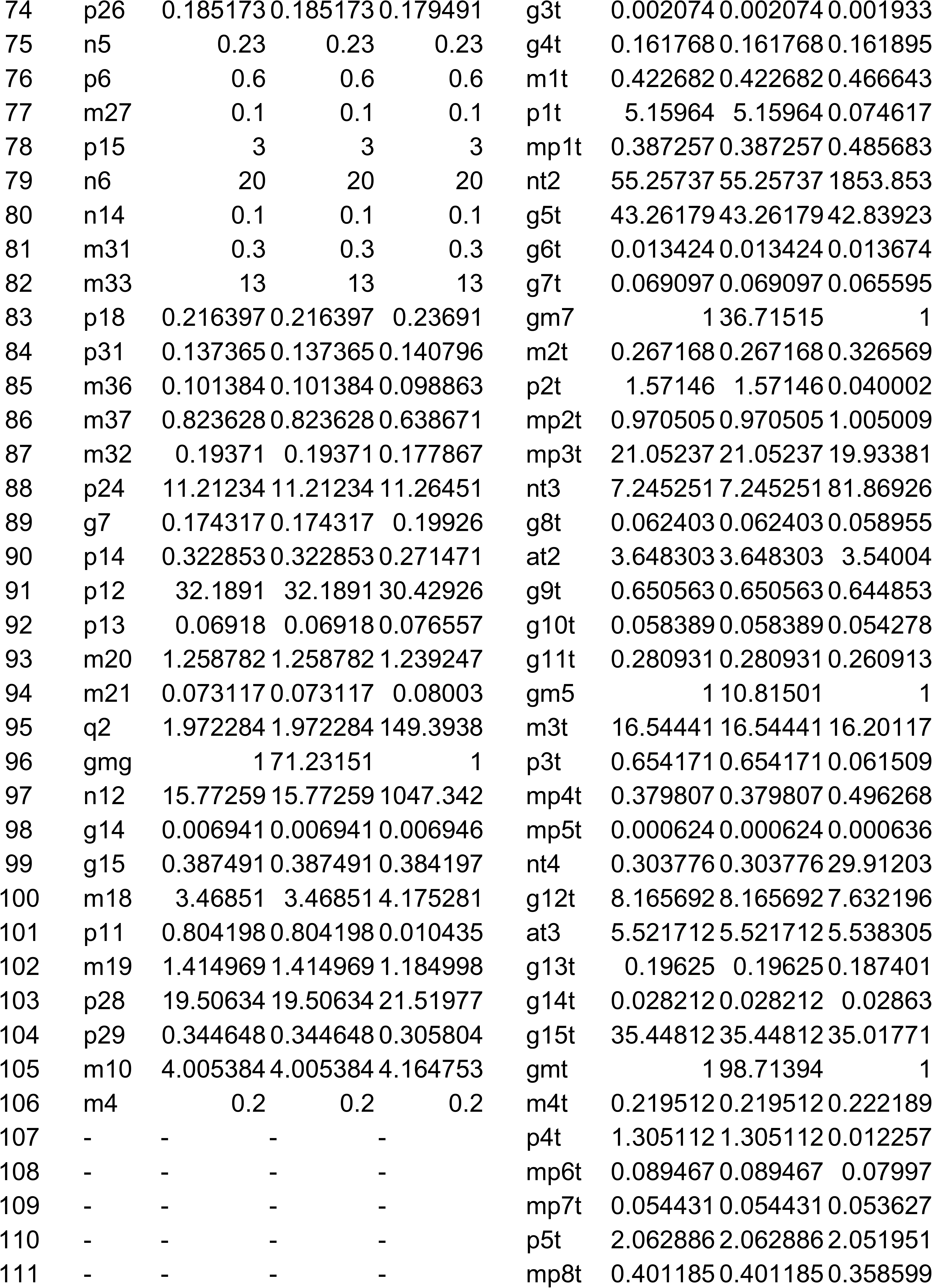
Parameter values for the models developed in this work. Hill coefficients were fixed to 2 and are not listed here but are included in the parameter total in Supplementary Table 2. The eight RNA scaling factors (named gmX) used to match the RNA data in absolute units are listed here but are not included in the parameter total. The order in the table corresponds to the order of parameters in the equations (see Supplementary Information).

## Supplementary Figure Legends

**Supplementary Figure 1.**
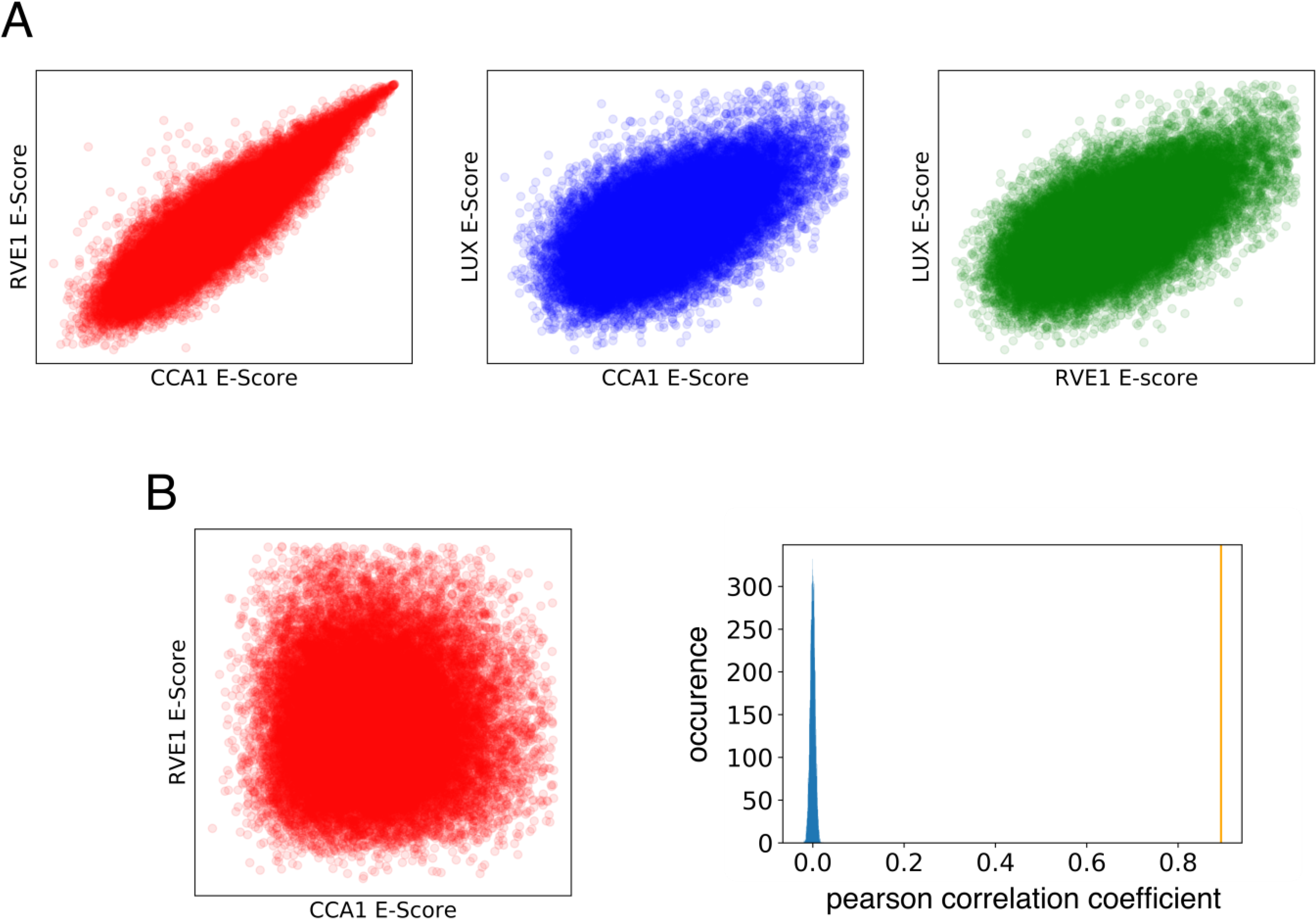
Similarity of DNA binding motifs between three clock proteins. E-scores for binding to test DNA sequences from protein binding microarrays (Franco-Zorrilla *et al*. 2014) are plotted for the clock proteins indicated. a) Pairwise comparisons of E-score between CCA1/RVE1, CCA1/LUX and RVE1/LUX show that the CCA1 and RVE1 binding site preferences are more similar than other comparisons. b) Bootstrap analysis to assess the significance of the correlation coefficients. 1,000 data permutations were tested (examples in red, left panel), resulting in the blue distribution of Pearson correlation coefficients (right panel). The observed correlation coefficient of the CCA1/RVE1 comparison in panel a) is also marked (yellow line).

**Supplementary Figure 2.**
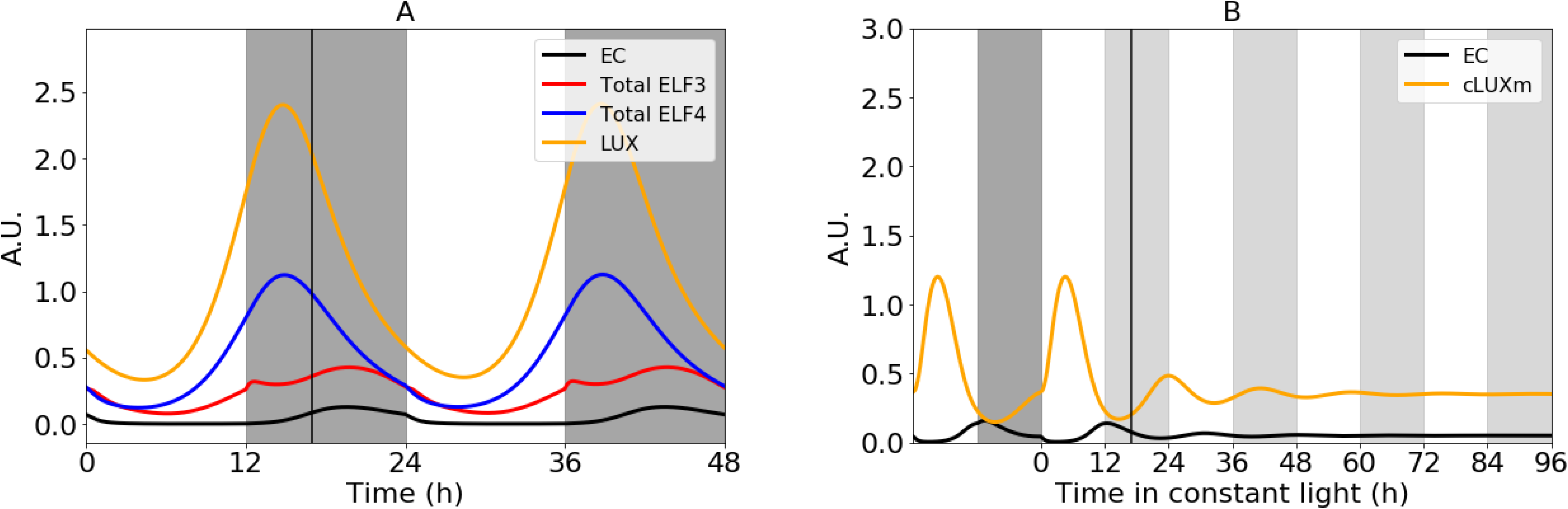
The Evening Complex (EC) in the P2011 model and damped oscillation in the *lhy/cca1* double mutant. a) Levels of the EC and its constituent proteins, ELF3, ELF4 and LUX, were simulated using the P2011 model under light:dark cycles (white: dark grey shading). EC levels peak after 17h (line), well after the ELF4 and LUX protein levels peak. b) accumulation of the *LUX* RNA oscillates nearly in antiphase to its auto-repressor the EC, in a simulation of the *lhy/cca1* mutant under L:D cycles (white: dark grey shading) before transfer to constant light (white: light grey) at time 0.

**Supplementary Figure 3.**
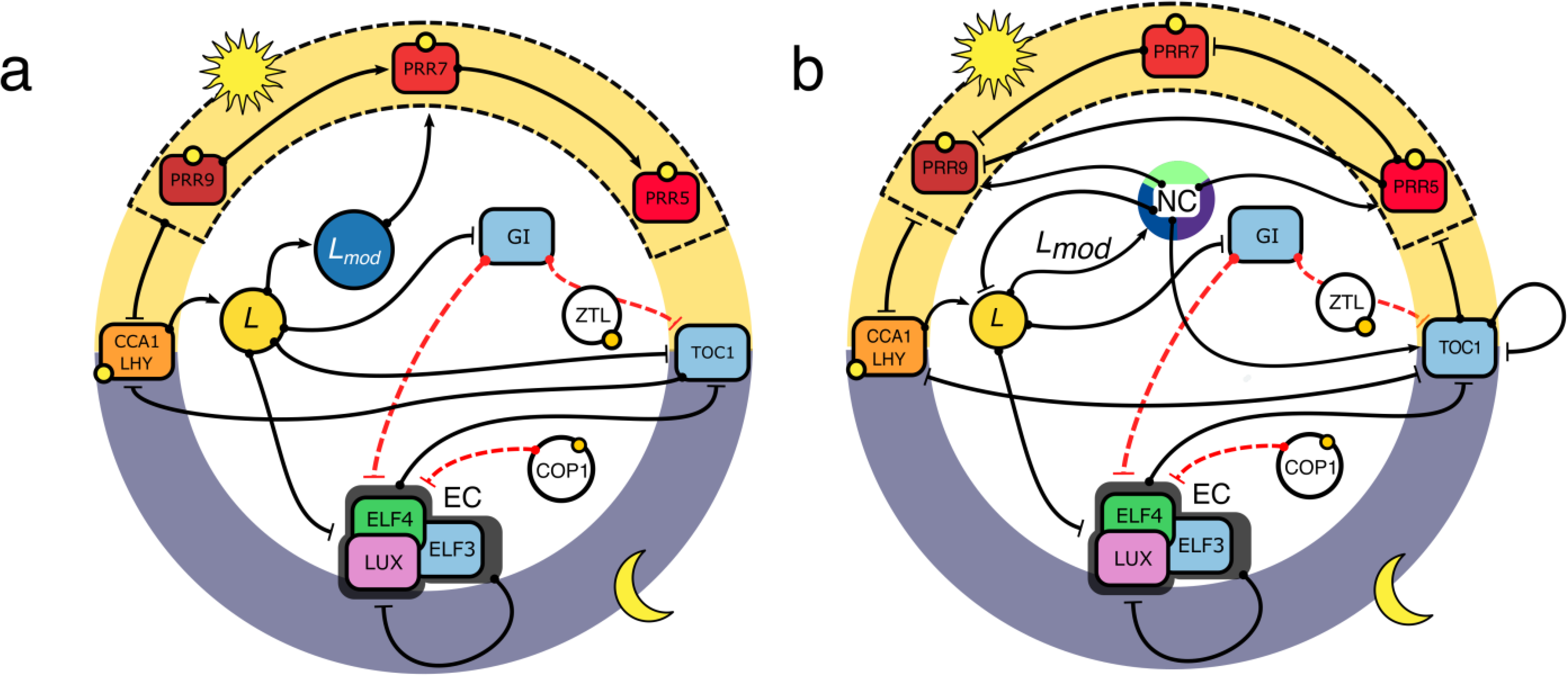
Simplified depiction of the U2019 (a) and U2020 (b) models, using conventions as in Figures 1a, 2a. Arrows, activation; blunt lines (T), repression. Solid edges, transcriptional regulation; dashed red lines, posttranslational regulation. Yellow circles, light regulation. The diel cycle is represented by the yellow (day) - grey (night) circle. The equivalent regulation of *CCA1/LHY* by each PRR protein is indicated the dashed black box. The Noon Complex (NC) proposed to function similarly to *Lmod* is shown as a pie, potentially formed from RVE, LNK and at least a third unknown component.

**Supplementary figure 4.**
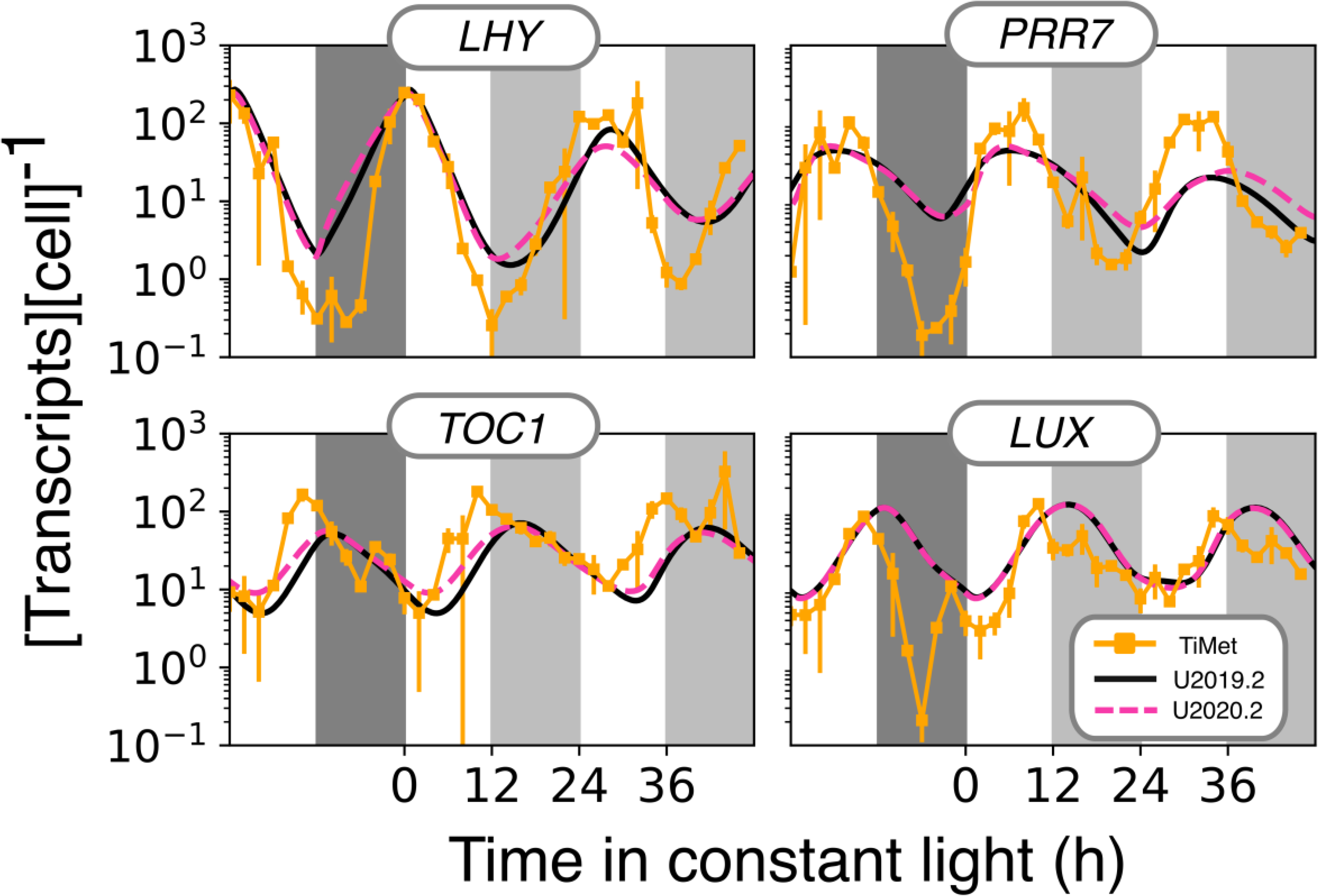
RNA levels in models U2019.2 and U2020.2 were rescaled to match TiMet data. RNA levels for four transcripts in Col-0 WT plants are shown, compared to simulation results from U2019.2 (solid black line) and U2020.2 (dashed red line). The model behaviour is close to U2019.1 and U2020.1 (Figure 3c) but matches the RNA levels of the TiMet data and hence the scaling of models U2019.3 and U2020.3 (Figure 5). The models were entrained for 10 days in L:D cycles (white:dark grey bars) before transfer to constant light (white:light grey) at time 0. Errors bars for experimental data are 1 S.D., n=2.

**Supplementary Figure 5.**
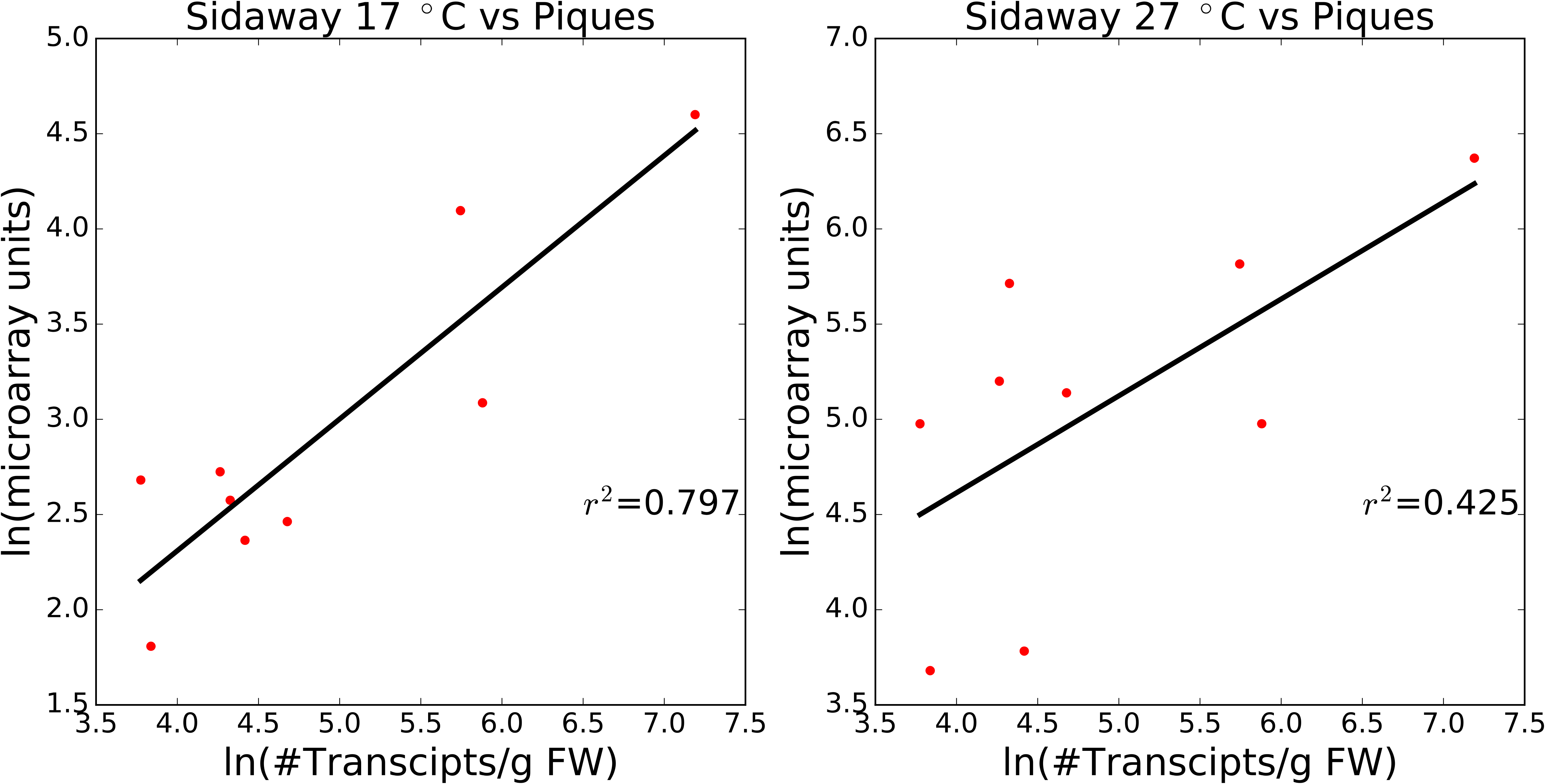
Condition-dependent correlation in RNA levels between the Piques data and Sidaway-Lee data. The linear regression for the ln-transformed levels of selected transcripts from the Piques data (in units of ln (microarray units)) against their levels in the Sidaway-Lee data obtained at 17C (left panel; Figure 6c; in units of ln(transcripts per gram fresh weight)) shows a higher correlation coefficient than the same Piques data against the Sidaway-Lee data obtained at 27C (right panel).

## Supplementary Information

### Comparison of the detailed models P2011 and F2014

The P2011 and F2014 models represent *LHY* and *CCA1* differently. P2011 uses a single, combined variable *CCA1/LHY* that activates *PRR9* in the morning and represses *TOC1, GI*, *ELF3*, *ELF4* and *LUX*. F2014 has specific variables for *LHY* and *CCA1*, which are distinguished by the autoregulation of *CCA1* by its cognate protein. CCA1 and LHY proteins in F2014 act as repressors of the *PRRs* and *GI, ELF4, ELF3, LUX* and its homologue *BROTHER OF LUX ARHYTHMO (BOA)/ NOX,* which was not represented in the P2011 model.

The temporal wave of PRR proteins represses the expression of *CCA1* and *LHY*. The earliest-expressed gene *PRR9* ends the dawn peak of *LHY* and *CCA1*, while the latest-expressed *TOC1* controls their rise in expression in the middle of the following night. In P2011 the wave of *PRR* expression is modelled through a “forwards activation” cascade of *CCA1/LHY*-> *PRR9*-> PRR7-> PRR5, and repression of only *TOC1* by CCA1/LHY, because construction of the model preceded evidence of CCA1/LHY’s widespread repressive functions. Matching the observed *PRR* dynamics also required a transcriptional activator (Pokhilko *et al*. 2010). F2014 achieves similar temporal *PRR* expression by a different mechanism, in which the PRR proteins function in a “backwards repression” cascade: later-expressed *PRR* genes repress the promoters of earlier-expressed genes. The F2014 model also introduced a variable RVE8 (discussed below) that represents the transcription-activating function of some members of the *REVEILLE / LHY-CCA1-like* family of transcription factors, which are dawn-expressed genes homologous to *LHY* and *CCA1*. Family members notably RVE 8 /LCL 5 have been shown to interact with a family of transcriptional co-activators termed NIGHT LIGHT-INDUCIBLE AND CLOCK REGULATED GENES, the LNK1-LNK4 family (Rugnone *et al*. 2013), which were not represented in either model.

The evening-expressed genes *LUX, ELF4* and *ELF3* together produce the hetero-trimeric protein Evening Complex (EC) (Nusinow *et al*. 2011). The P2011 model simulates this stepwise. ELF3 and ELF4 form a dimer, which then binds to LUX resulting in the formation of the EC. In the F2014 model, the EC representation is more phenomenological, using an *ad hoc* algebraic equation that also includes BOA/NOX. In both models, the EC binds to the promoters of *PRR9, TOC1/PRR1, LUX, ELF4* and *GI* and represses their expression.

Both models explicitly represent the dynamics of light-regulated proteins, among the major mechanisms of light input. These include the CONSTITUTIVE PHOTOMORPHOGENESIS 1 (COP1) system and the interaction of the F-box protein ZEITELUPE (ZTL) with GI and PRR proteins. COP1 protein dynamics are identical in both models, with minor differences in its effects on EC formation due to the EC’s differing representation in the models. The regulation of *GI* transcription and ZTL:GI interactions are represented with slightly more complexity in the F2014 than the P2011 model, reflecting later publications. In P2011, light destabilises the proteins PRR9, PRR7 and PRR5, whereas in F2014 only PRR7 stability is controlled in this way. Light-regulated transcription is represented in both models, through continuous effects of light and through an acute, transient response at dawn. The latter is represented using the hypothetical protein P (J. C. W. Locke *et al*. 2005), which accumulates in dark intervals, is rapidly degraded upon illumination and was recently identified with phytochrome A (PhyA) (Seaton *et al*. 2018).

### Equations for models U2019 and U2020

Equations generated from the SBML files using COPASI software are detailed on the following pages, for U2019 followed by U2020. The parameter *Vcell* is automatically added by the software but is not used in this model; it has a value of 1. The equation used to generate the light:dark cycle is omitted. The SBML model files incorporate the standardised Input Signal Step Function, which provides a tunable light:dark cycle based on hyperbolic tangent functions that are mathematically continuous (Adams et al. 2012), and a transition from LD to LL encoded in an SBML Event. An alternative step function could be substituted, with little effect on model dynamics.

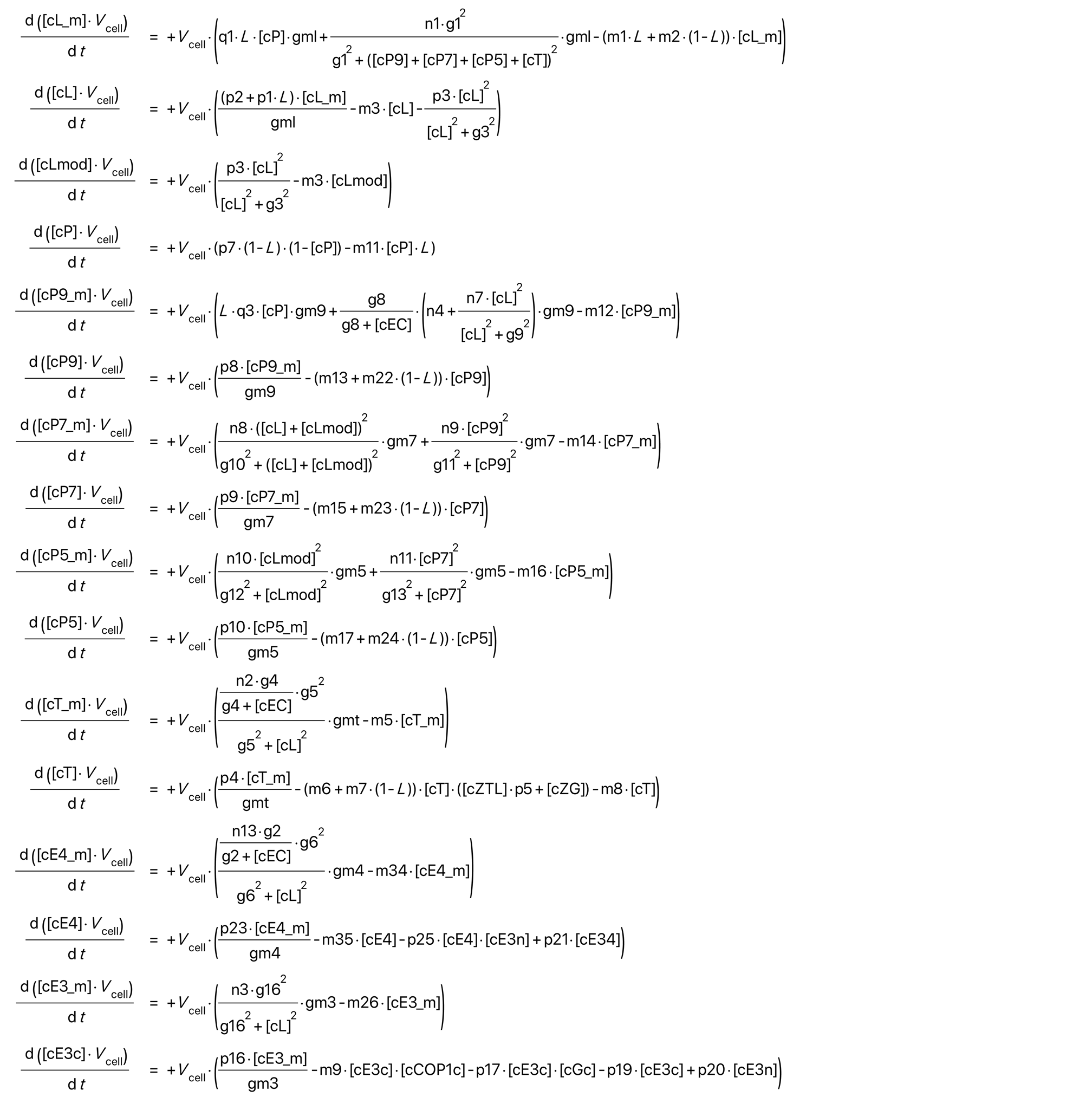

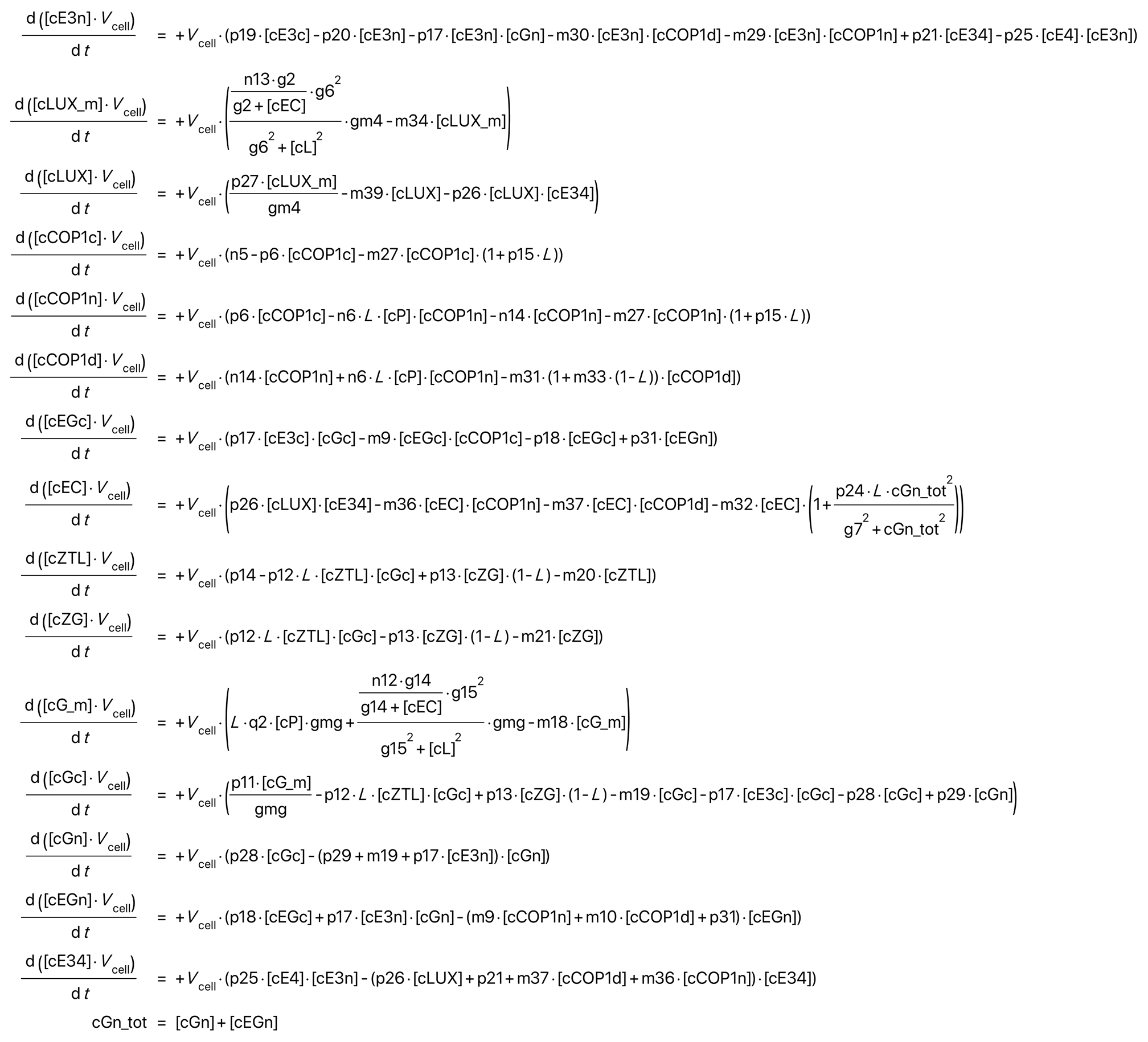

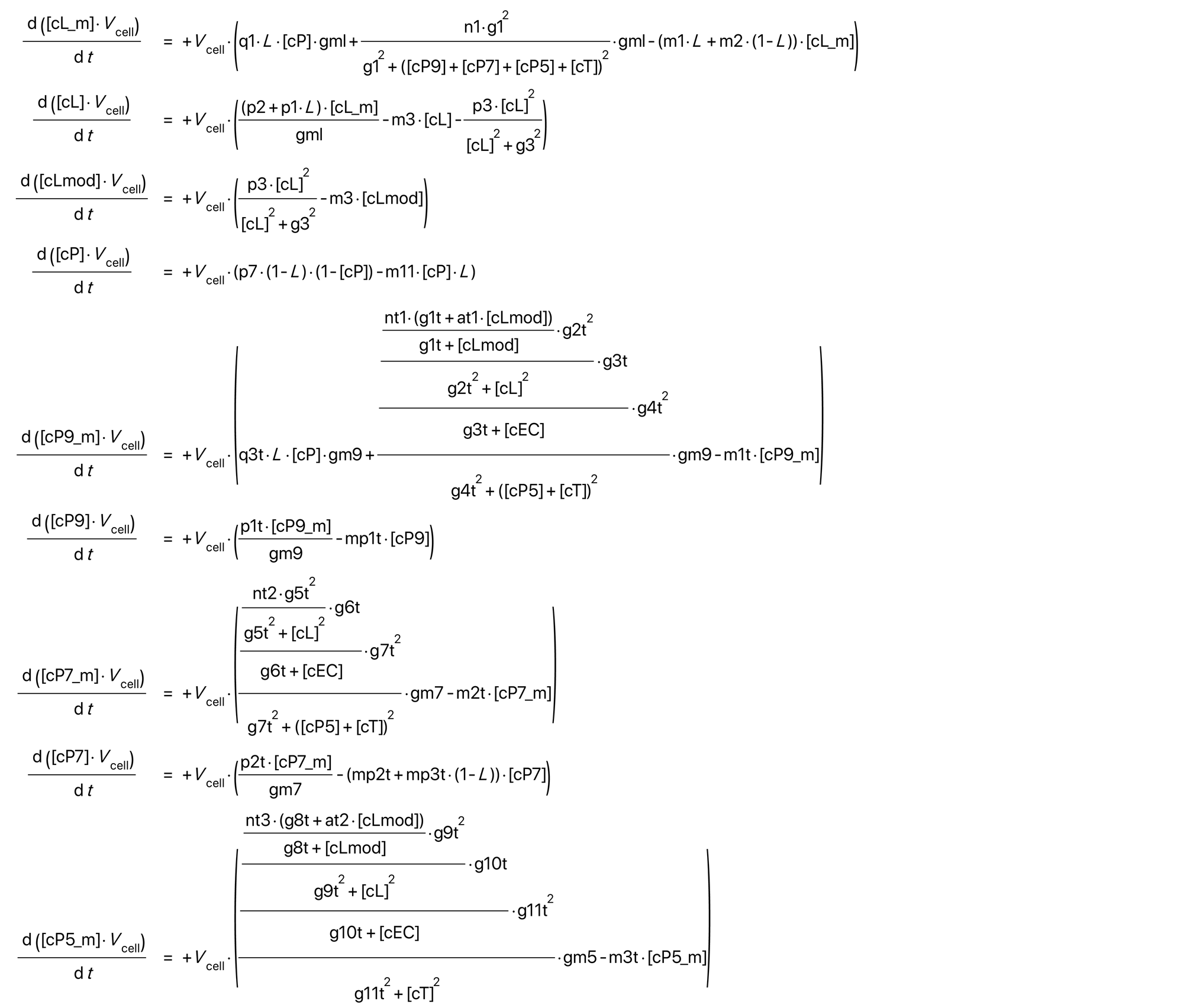

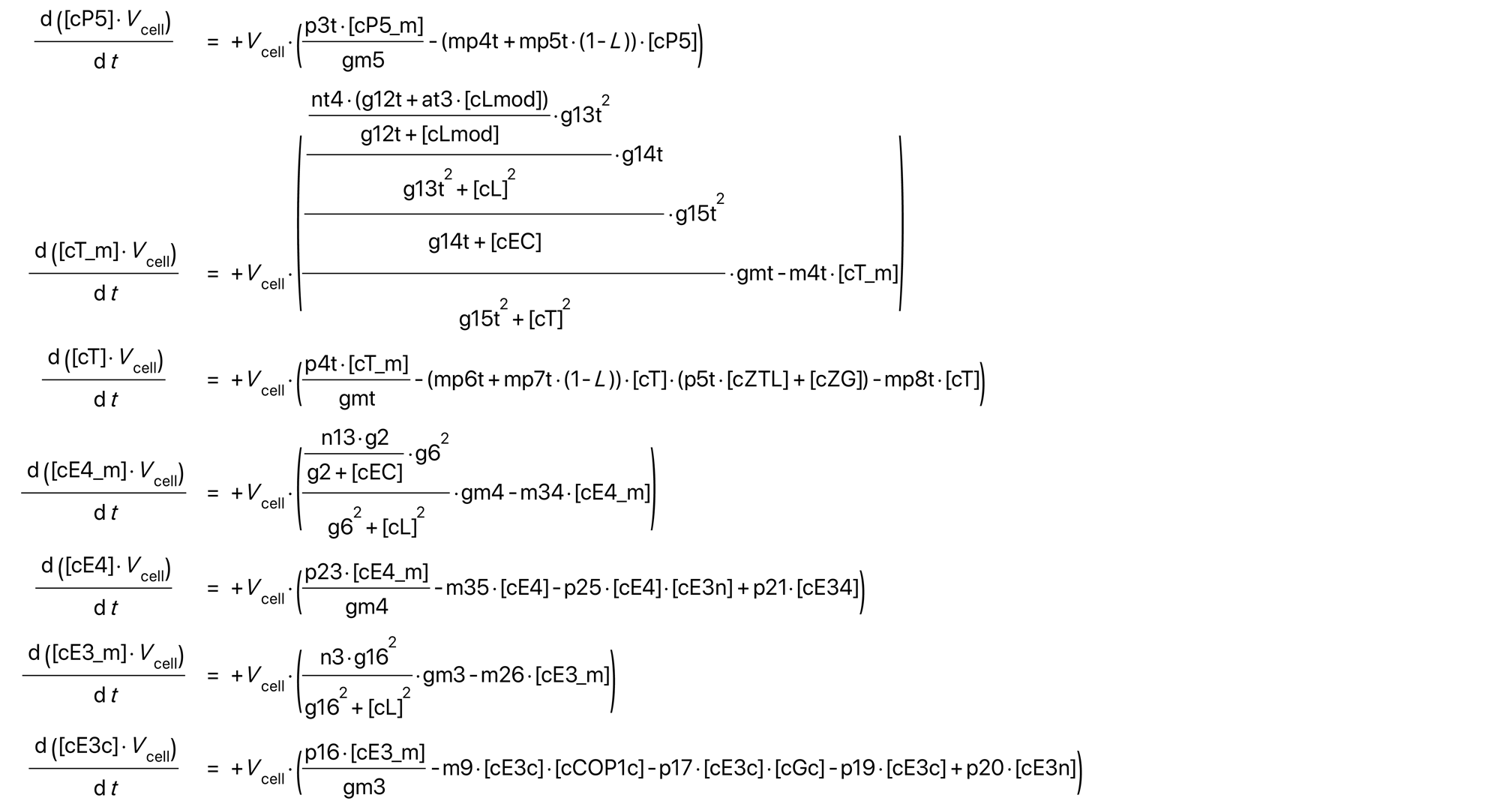

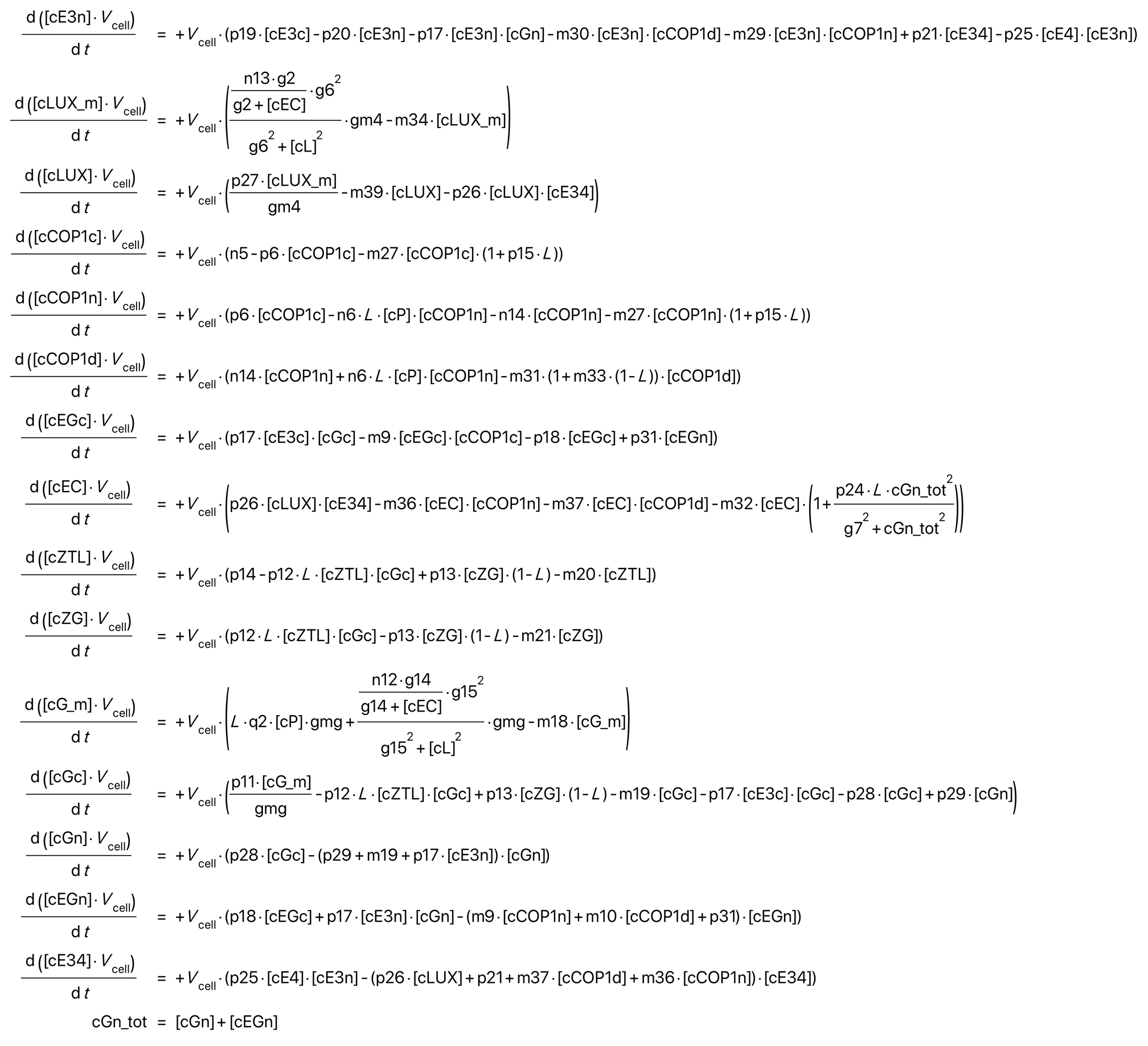

